# Integrative analysis with expanded DNA methylation data reveals common key regulators and pathways in cancers

**DOI:** 10.1101/491845

**Authors:** Shicai Fan, Jianxiong Tang, Nan Li, Ying Zhao, Rizi Ai, Kai Zhang, Mengchi Wang, Wei Du, Wei Wang

## Abstract

The integration of genomic and DNA methylation data has been demonstrated as a powerful strategy in understanding cancer mechanisms and identifying therapeutic targets. The TCGA consortium has mapped DNA methylation in thousands of cancer samples using Illumina Infinium Human Methylation 450K BeadChip (Illumina 450K array) that only covers about 1.5% of CpGs in the human genome. Therefore, increasing the coverage of the DNA methylome would significantly leverage the usage of the TCGA data. Here, we present a new model called EAGLING that can expand the Illumina 450K array data 18 times to cover about 30% of the CpGs in the human genome. We applied it to analyze 13 cancers in TCGA. By integrating the expanded methylation, gene expression and somatic mutation data, we identified the genes showing differential patterns in each of the 13 cancers. Many of the triple-evidenced genes identified in the majority of the cancers are biomarkers or potential biomarkers. Pan-cancer analysis also revealed the pathways in which the triple-evidenced genes are enriched, which include well known ones as well as new ones such as axonal guidance signaling pathway and pathways related to inflammatory processing or inflammation response. Triple-evidenced genes, particularly TNXB, RRM2, CELSR3, SLC16A3, FANCI, MMP9, MMP11, SIK1, TRIM59, showed superior predictive power in both tumor diagnosis and prognosis. These results have demonstrated that the integrative analysis using the expanded methylation data is powerful in identifying critical genes/pathways that may serve as new therapeutic targets.

## Introduction

The Cancer Genome Atlas (TCGA, https://tcga-data.nci.nih.gov/tcga/) has profiled the genomic and epigenomic variations of thousands of samples for several dozens of cancers^1^. These multiomics data include genetic variation, gene expression and DNA methylation that provide an invaluable resource for understanding the cancer mechanisms and identifying new therapeutic targets. A limitation of the TCGA DNA methylation data is that it was generated using Illumina Infinium Human Methylation 450K BeadChip (referred to as Illumina 450K array hereinafter), which only covers about 1.5% of the CpGs in the human genome. This poor coverage restricts epigenomic analysis and many differentially modified loci are likely missed. While Whole Genome Bisulfite Sequencing (WGBS) and other technologies are available to measure DNA methylation with much higher coverage, it is unlikely to repeat the DNA methylation analysis in the large number of TCGA samples considering the expense and effort in the near future. Therefore, there is an urgent need to develop new analysis strategy to better use these data.

Previously, we developed a method to expand the Illumina 450K array data by considering sequence features and local methylation profile in the neighboring CpGs^2, 3^. Despite the promising results provided by these methods, their speed is slow and applying them to expand the thousands of TCGA data is infeasible. Here, we present an improved model called EAGLING (Expanding the 450K methylation Array with neiGhboring methylation value and Local methylation profilING) with a more than 10 times faster speed compared to our previous methods. Furthermore, the location distribution of the expanded CpG sites is less biased towards CpG rich regions, and the hyper/hypo-methylated ratio is also more similar to the ratio from the WGBS data. Importantly, the coverage of CpGs is significantly increased from about 1.5% of all CpGs in the human genome in the original Illumina 450K data to about 30% after expansion.

This new model allows integrative analysis of genetic variation, gene expression and expanded DNA methylation to identify genes and pathways that are important for diagnosis and therapeutic treatment. We identified the triple-evidenced genes in each of the 13 TCGA cancers that have sufficient samples. The triple-evidenced genes represent the genes that are differentially methylated, differentially expressed and associated with somatic mutation. We found that the triple-evidenced genes shared by a majority of the 13 cancers include many previously identified biomarkers or therapeutic targets^4-7^. These triple-evidenced genes are enriched in numerous pathways, suggesting possible new targets for therapeutics. Importantly, these triple-evidenced genes can discriminate the cancer from normal samples and predict survival. In particular, nine genes, TNXB, RRM2, CELSR3, SLC16A3, FANCI, MMP9, MMP11, SIK1, TRIM59 are important in both cancer diagnosis and prognosis; note that FANCI and SIK1 would be missed if using the original Illumina 450K data. The EAGLING model and all of the triple-evidenced genes are available at http://114.55.236.67:8013/Integrative_Analysis/home.

## Results

We propose here an integrative analysis strategy to identify key regulators and pathways in cancers from the TCGA data. By comparing gene expression, genetic variation and DNA methylation data between normal and cancer samples, we extracted the triple-evidenced genes for 13 cancers and analyzed the characteristics of these genes (Figure 1A).

**Figure 1.**
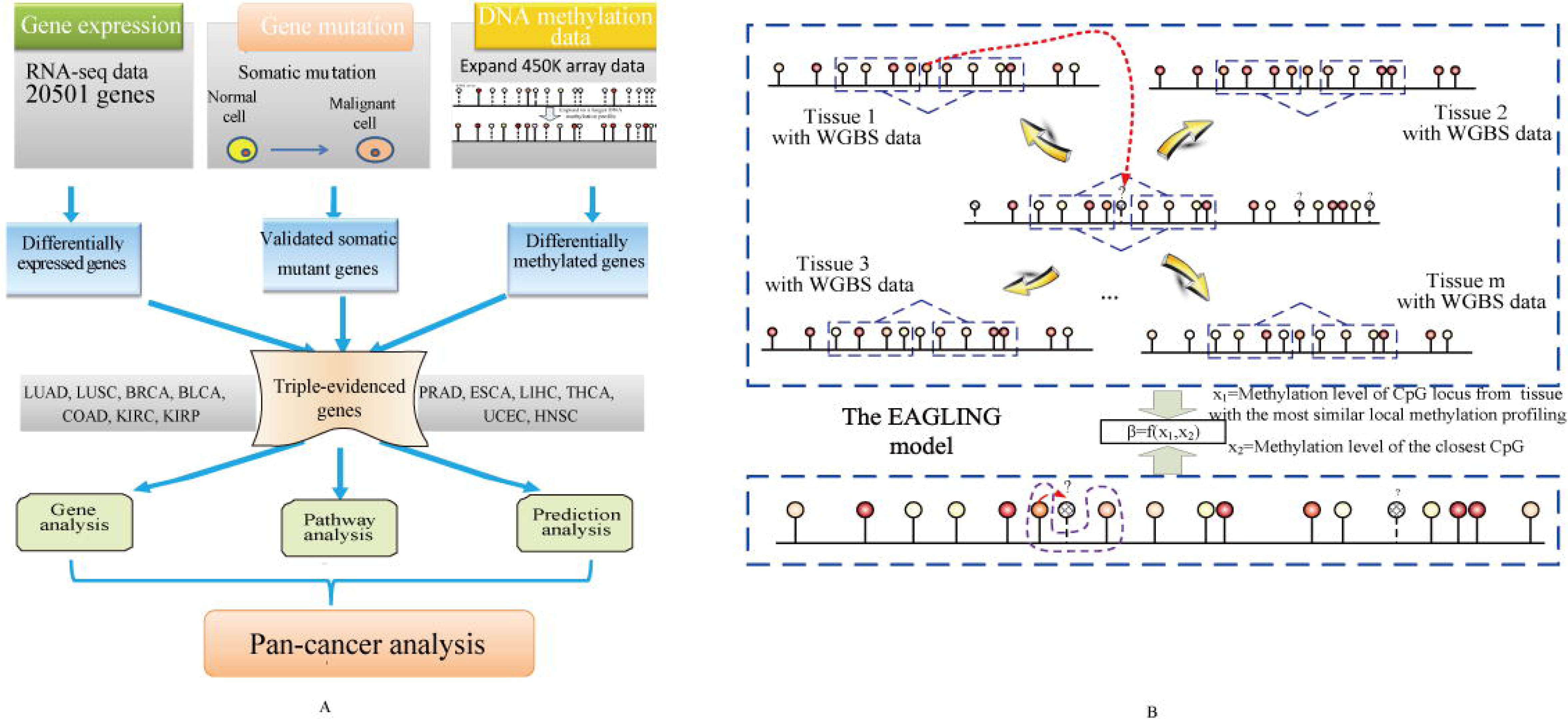
The workflow of the integrative analysis and the EAGLING model. (A) The multiomics data of 13 cancers from TCGA were used to identify the genes that are differentially expressed and differentially methylated and also contain somatic mutations (i.e. triple-evidenced genes) in each cancer. Pan-cancer analysis revealed that the triple-evidenced genes shared by a majority of the 13 cancers include many previously identified biomarkers or therapeutic targets. (B) In the model construction, two features are used to build the logistic regression model: the methylation level of the closest CpG based on 450K array and the methylation value from the WGBS of the corresponding CpG from the tissue that has the most similar local methylation profile with the site to be predicted.

### The EAGLING model expands the 450K array methylation data 18 times

In order to expand the Illumina 450K array DNA methylation data, we previously developed prediction models based on local methylation patterns and sequence features. In this work, we proposed a new model dubbed as EAGLING that has two steps to predict methylation levels of CpGs based on the Illumina 450K array data. First, it finds a WGBS methylome that shares the most similar local methylation profile around the CpG L under consideration, and the methylation value of the CpG in the selected WGBS methylome is taken as an input feature; second, the methylation value of the closest CpG from Illumina 450K array is taken as another input feature. A logistic regression model was built on these two features to predict the methylation level at L (Figure 1B, see details in Materials and Methods). Note that this procedure is repeated for each CpG so that different CpGs may be predicted from different WGBS methylomes. Thirty-three tissues/cell lines in which both 450K array and WGBS data were available were used to optimize the parameters. There are three major improvements over our previous models^2, 3^. First, DNA sequence features are not included in EAGLING, which significantly improves the speed without deteriorating the performance; second, the methylation value of the closest CpG is used because of its higher precision compared to the weighted neighbor CpGs used in our previous models^2, 3^; third, more training data are included (33 versus 14 tissues/cell lines), which is expected to improve the model.

We have searched for the optimal number of CpGs to represent the local methylation pattern in step 1, which is crucial to identify the WGBS methylome from which to take the methylation level for the CpG under consideration. We performed the leave-one-tissue-out cross validations using 1-10 neighbor CpGs and 5Kbp-50Kbp flanking regions (Figure 2A). The flanking regions confine the CpGs we considered. Only the CpGs with the required number of neighbor CpGs in the specified flanking regions were included for expansion because their local methylation profiles could be accurately represented. We chose 4 CpGs each on upstream and downstream sides to represent the local methylation profile as there was no performance improvement by including more than 4 flanking CpGs and 30Kbp for the flanking region size considering the balance between satisfactory performance and genome coverage (Figure 2B). Using these parameters, the leave-one-tissue-out cross validations achieved superior performance (Figure 2C). To show the impact of the training size on the model performance, we trained the EAGLING model using 14 (the training sample size for our previous model in reference 3) and 23 WGBS data sets (the 14 WGBS data plus another 9 randomly chosen WGBS data) separately. We compared their prediction results on another 10 WGBS samples not included in the training set. We repeated this cross validation 10 times and the results are shown in Figure S1a. The EAGLING model trained using 23 WGBS data showed improved correlation coefficient (0.8441), concordance (0.8532), accuracy (89.65%) and AUC (0.8645) compared to those trained using 14 WGBS data (0.8321, 0.8375, 87.11% and 0.8595). Importantly, not including DNA sequence features in EAGLING does not impair the prediction performance while removing the time consuming step of considering sequence features in the previous model (Figure S1b) to achieve a 10 times faster speed.

**Figure 2.**
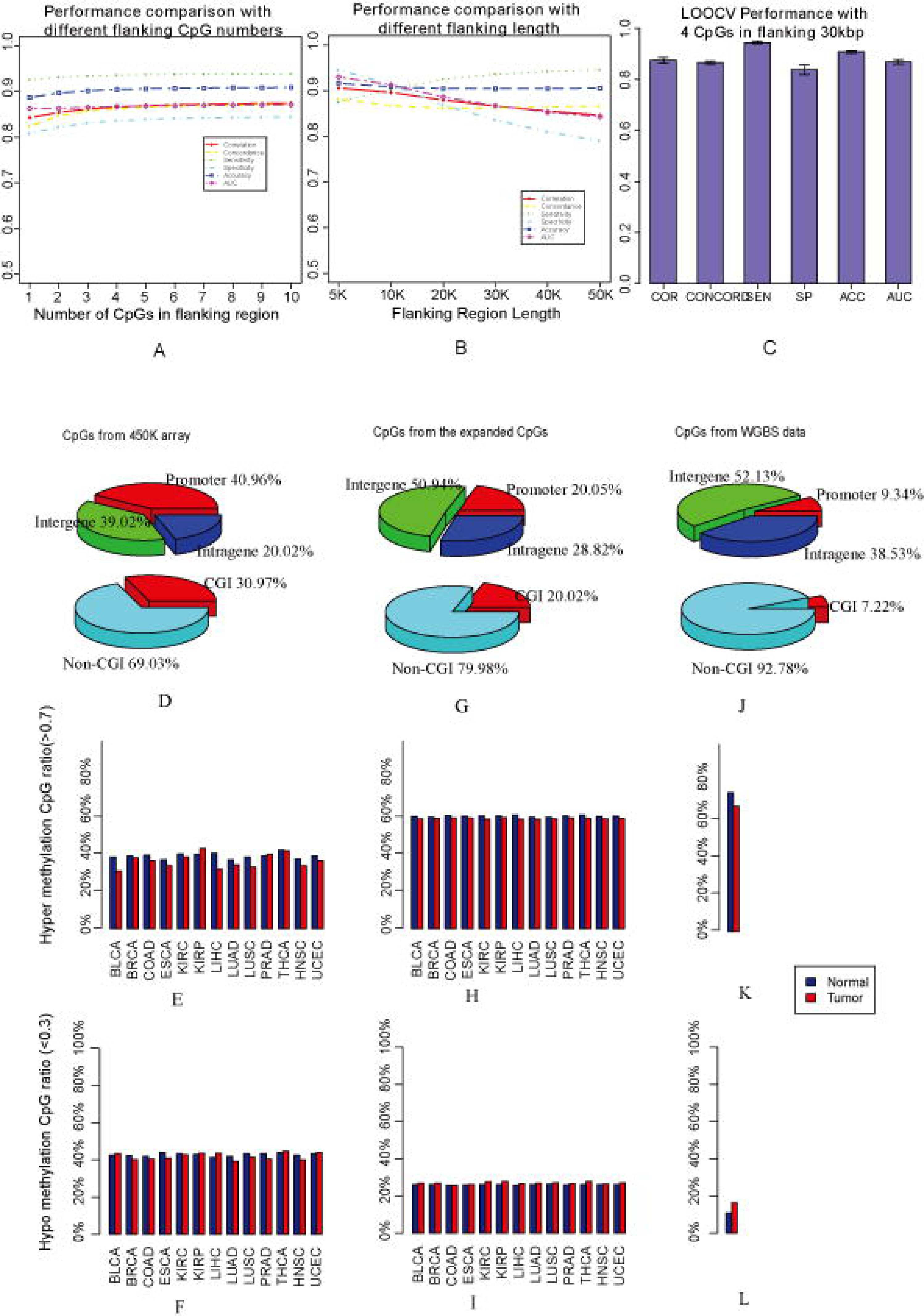
The performance of EAGLING model and the expanded methylation profile. (A) and (B) The leave-one-tissue-out cross validations with different flanking CpG numbers and sizes of the flanking regions. (C) The performance with 4 CpGs in the flanking 30kbp regions. Pearson correlation coefficient (COR), Concord (CONCORD, the percent of CpGs with a methylation proportion difference less than 0.25 ^60^), Sensitivity (SE), Specificity (SP), Accuracy (ACC), and AUC (Area Under ROC Curve) are used as the metrics to assess the performance. (D), (E) and (F) are the CpG location, hyper/hypo methylation ratio in tumor/normal samples from the Illumina 450K array, respectively; (G), (H) and (I) are the CpG location, hyper/hypo methylation ratio from the expanded data, respectively; (J), (K) and (L) are the CpG location, hyper/hypo methylation ratio from the WGBS data, respectively.

Furthermore, we applied our EAGLING model on 450K array data of K562 and HepG2, two independent cancer cell lines from the ENCODE project. The predicted methylation levels were well correlated with the WGBS data in the same cell lines: the correlation, accuracy and AUC on K562 and HepG2 were 0.84, 84.13%, 0.84 and 0.91, 92.27%, 0.87, respectively. The correlation, accuracy and AUC in K562 and HepG2 using our previous model^3^ were 0.82, 81.13%, 0.80 and 0.89, 90.13%, 0.82, respectively. The superior performance further validated the EAGLING model.

### Expanding the Illumina 450K array data in the TCGA samples

Using the EAGLING model trained on the 33 tissues/cell lines (see details in Table S1), we expanded the Illumina 450K methylation array data in 13 cancers from TCGA (LUAD, LUSC, BRCA, BLCA, COAD, KIRC, KIRP, PRAD, ESCA, LIHC, THCA, UCEC and HNSC) that have at least 10 normal samples of Illumina 450K array and RNA-seq data. The expanded data increased the coverage of CpGs to 18.9 times (about 30% of CpGs in the human genome). Particularly, the intergenic coverage was significantly increased from 39.02% in 450K array to 50.94% in the expanded data and the non-CpG island coverage also increased from 69.03% to 79.98%, which is important to identify functional enhancers (Figure 2D and 2G). The location distribution of the expanded data is much closer to that of all the CpGs in human genome (Figure 2J) than the original 450K array. Furthermore, we identified the hyper-methylation (>0.7) and hypo-methylation (<0.3) CpGs from the original 450K array data and calculated their percentages among all the CpGs for the tumor and normal samples of the 13 cancers (Figure 2D and 2E). Obviously, the ratio distributions of the expanded CpGs are much closer to those of the WGBS data (Figure 2K and 2L), indicating that the analysis results based on the expanded methylation data would be less biased compared to the results from the 450K array data.

### Identification of triple-evidenced genes

Using the expanded methylation data in the 13 cancers, we identified the differentially methylated genes (DMGs) between the tumor and normal samples (see Methods for detail). We also identified the differentially expressed genes (DEGs) using the RNA-seq data and genes containing somatic mutations (see details in Methods and Figure S2). As an example, the overlap between DMGs, DEGs and genes with somatic mutation of LUSC is shown in Figure 3A. In the 13 cancers, the number of triple-evidenced genes ranges from 396 in PRAD to 1438 in LUSC (Figure S2).

**Figure 3.**
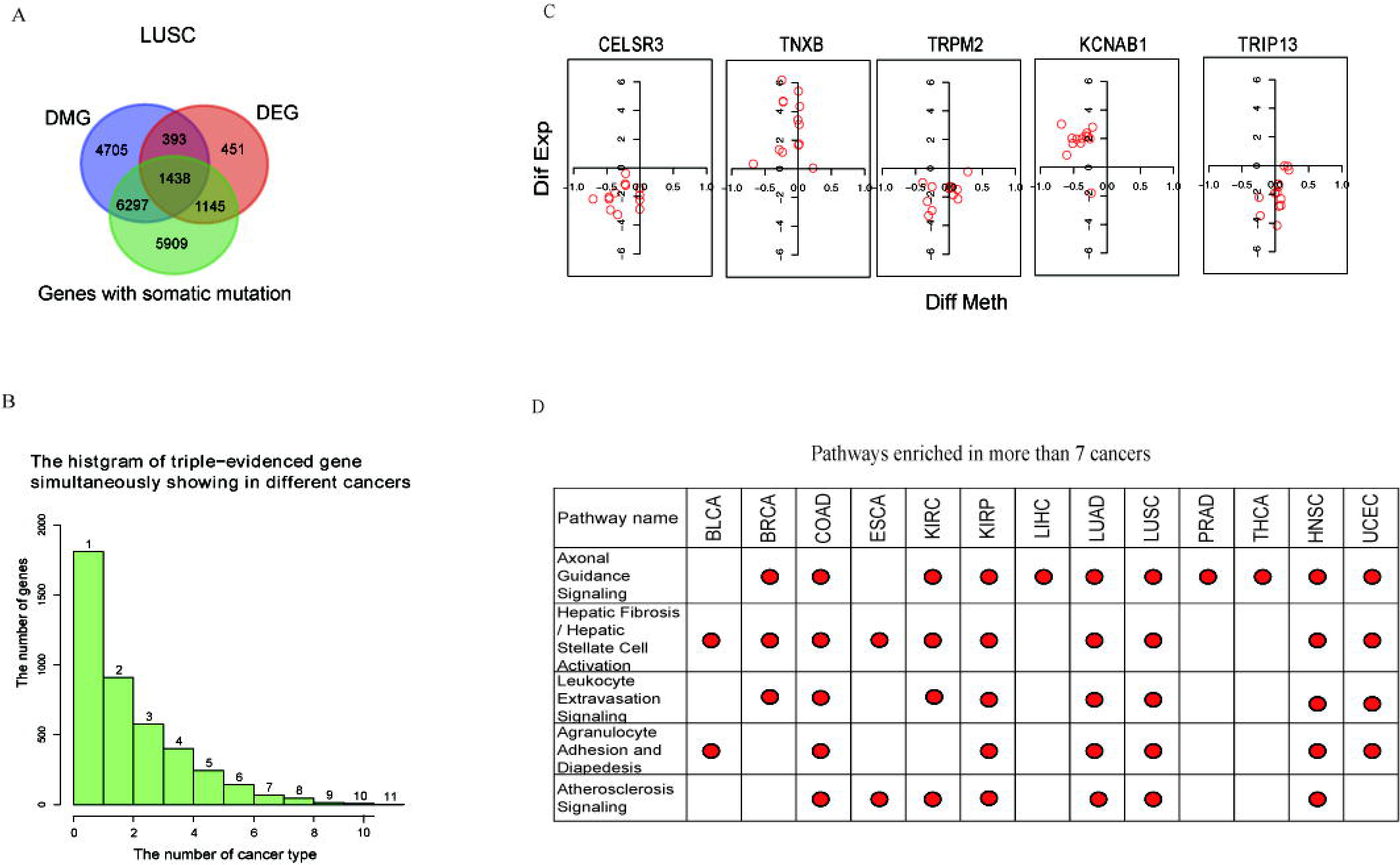
Analysis of triple-evidenced genes and enriched pathways. (A) Genes supported by one, two or three evidences in LUSC. (B) The number of triple-evidenced genes shared between the 13 cancers. (C) The differential methylation and expression levels of the top 5 triple-evidenced genes (difference=normal value - tumor value) that are most common in the 13 cancers. X axis is the difference of methylation, y axis is the log2(RPKM ratio) to represent the gene expression difference. One red circle in each graph represents a cancer type. (D)The top 5 pathways enriched in the 13 cancers, the red dot indicates the cancer type in which a pathway is enriched.

Only a small portion of the triple-evidenced genes were found in more than 6 cancers (Figure 3B). The top 5 triple-evidenced genes found most often in the 13 cancers are listed in Table S2. They were CELSR3, TNXB, TRPM2, KCNAB1 and TRIP13 that were identified as triple-evidenced genes in 11, 11, 11,10 and 10 types of cancers, respectively. We first examined the difference of their methylation and expression levels between tumor and normal samples (Figure 3C) (difference = normal value - tumor value). CELSR3, TRPM2 and TRIP13 are over-expressed in all the 13 cancers, TNXB is under-expressed in all the 13 cancers, KCNAB1 is over-expressed in KIRC but under-expressed in the other 12 cancers. These genes show abnormal but consistent expression patterns across different cancers. The methylation level does not show clear trend though, indicating that the relationship between gene expression and their promoter methylation is complex, which is consistent with the previous studies^8, 9^.

Four of the 5 genes have been reported as biomarkers, potential biomarkers or therapeutic targets. CELSR3 was found to be highly expressed in ovarian cancer^4^, and hypermethylated in primary oral squamous cell carcinoma, and might be used as a biomarker in OSCC prognostication^10^ and small intestinal neuroendocrine tumor^5^. TNXB was reported to be important for the tumorigenesis of lung adenocarcinoma^6^, and was validated as a promising biomarker for early metastasis of breast cancer^7^.TRPM2 was reported to be a potential target of the selective treatment of prostate cancer^11^ and was suggested to be a potential therapeutic target in breast cancer^12^. TRIP13 promoted early steps of the DNA double-strand break repair and its presence was associated with progression in prostate cancer and squamous cell carcinoma of the head and neck^13, 14^. For KCNAB1, there were few reports about its function in cancer, but it was downregulated in follicular carcinoma and could be combined with other genes for the classifier construction^15^.

As a comparison, we also identified the triple-evidenced genes using the Illumina 450K data (Figure S3). There are two advantages using the expanded methylation data. First, the triple-evidenced genes can be identified in more cancers. For example, the CELSR3 gene was found as the triple-evidenced gene in two cancers using the original 450K array data but in 11 cancers using the expanded data; second, consistently, more triple-evidenced genes can be identified in a particular cancer by the expanded data than the original 450K array data. For example, 5 genes (FANCI, RECQL4, TACC3, CLU and SIK1) were reported to function in different cancers ^10, 16-19^ but they could not be identified using the Illumina 450K array data in any of the 13 cancers; in contrast, all of them were found as triple-evidenced genes in more than 6 cancers using the expanded data (Table S3).

### Triple-evidenced genes are enriched in particular pathways

For each of the 13 cancers, we checked the enriched pathways among the triple-evidenced genes using Ingenuity Pathway Analysis (IPA) (with Benjamini-hochberg adjusted p-value < 0.05). Some enriched pathways are known to be important in cancer, such as MMPs (Inhibition of Matrix Metalloproteinases), VEGF Family Ligand-Receptor Interactions, Wnt pathway, NF-kB Signaling, MAPK Signaling. The top 5 pathways most often found in the 13 cancers are shown in Figure 3D.

Axonal guidance signaling, which belongs to Neurotransmitters and Other Nervous System/ Organismal Growth and Development Signaling, is enriched in 11 out of 13 cancers (Figure S4). Genes included in the pathway have been implicated in cancer cell growth, survival, invasion and angiogenesis^20^. It was also reported that pancreatic cancer genomes show aberrations in the axonal guidance pathway genes^21^. As an example, the triple-evidenced genes in LUSC overlapped with this pathway are marked in purple in Figure S4. The top genes shared in the 11 cancers on the pathway are marked with star shape. Some of them have been targeted by drugs to treat numerous cancers such as Marimastat for breast and lung cancer, and Dabrafenib for non small-cell lung cancer.

The other 4 enriched pathways are Hepatic Fibrosis/Hepatic Stellate Cell Activation, Leukocyte Extravasation Signaling, Agranulocyte Adhesion and Diapedesis and Atherosclerosis Signaling. All of them are involved in inflammatory process or response, and their top functions are in Cell-To-Cell Signaling and Interaction, Cellular Movement or Immune Cell Trafficking. The association between the development of cancer and inflammatory is well recognized^22^, and about 20% of human cancers are related to chronic inflammatory caused by infections, exposure to irritants or autoimmune disease^23, 24^. The details of the pathways (LUSC as an example) are shown in Figure S5-S8. Note that the number of cancers in which the enriched pathways was identified is significantly larger than that identified from the original 450K array data (Figure S9).

Several triple-evidenced genes (MMP9, MMP11, CXCL12, MYL9) appear in 3 out of the 5 significantly enriched pathways and in more than half of the 13 cancers. All of these genes have been reported associated with cancers. The promoter methylation of CXCL12 was acted as a prognostic biomarker in prostate cancer patients^25^ or sporadic breast cancer^26^. The low expression level of MYL9 is correlated with a significantly reduced median survival rate in colon cancer patients and might act as clinical biomarkers for the early diagnosis of colon cancer^27^. MMP9 and MMP11, both of which belong to Proteins of the matrix metalloproteinase (MMP) family, were reported as tumor biomarkers^28^ or associated with tumor survival^29^, and targeted by an inhibitor marimastat.

### Diagnostic power of the triple-evidenced genes

We then investigated whether the triple-evidenced genes are useful in distinguishing cancers from normal samples. We trained a random forest model with the selected triple-evidenced genes to discriminate the pooled cancer samples from the normal ones. Because there are a large number of triple evidenced genes in all the cancers, we chose those that appear in more than half of the 10 training cancer types as the candidates. Their associated gene expression and DNA methylation levels of individual CpGs in the promoters of the selected genes in the expanded data were input features; the somatic mutation information could not be included because the mutation information for each gene in every sample was not available. The model was constructed with a cross validation strategy by sampling 10 cancer samples as training data and the remaining 3 cancer samples as test data for 100 times. LASSO was applied to select features in constructing each random forest model. The features are presumably important if they were most often selected in the 100 cross validations. We list the 47 features selected in more than 50 times of the cross validations in Table S4, including expression of 13 genes and methylation levels of 34 CpGs. As shown in Figure 4A, most of the test tumor samples could be correctly predicted as cancers while about 75% of the test normal samples were predicted as normal samples. Obviously, using triple-evidenced genes derived from the expanded methylation data outperformed using the original 450K array data (t-test), particularly the specificity. The AUC of the cancer diagnosis with the triple-evidenced genes is about 0.85, which indicates that there are common differential gene expression and methylation features of the triple-evidenced genes that distinguish tumors from the normal samples.

**Figure 4.**
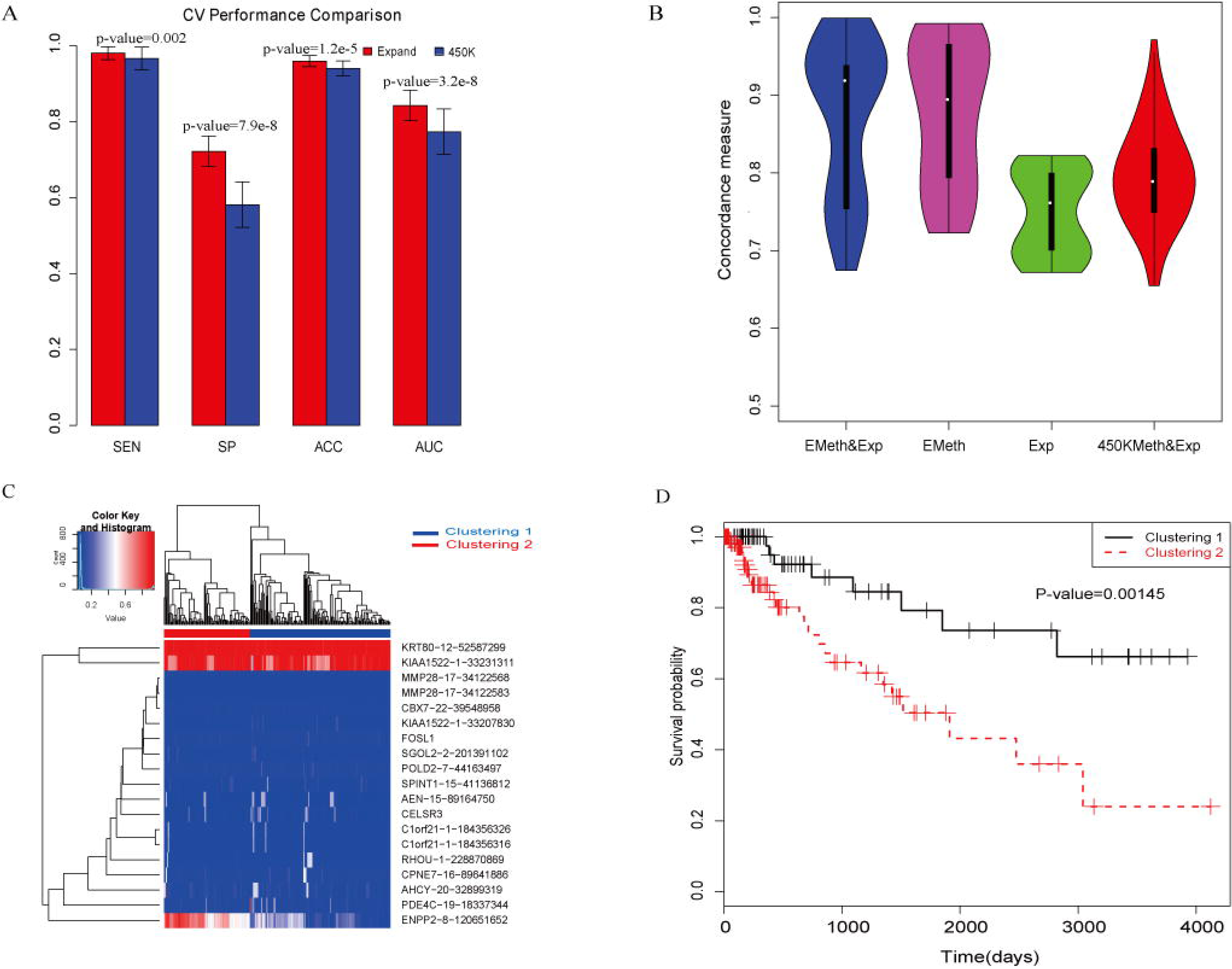
Diagnosis and prognosis analysis using the triple-evidenced genes. (A) Diagnostic analysis to distinguish cancers from normal samples using the triple-evidenced genes in 10-fold cross validations (t-test). Sensitivity (SE), Specificity (SP), Accuracy (ACC), and AUC (Area Under ROC Curve) are used as the metrics to assess the performance. (B) The boxplots of the concordances using expression data alone, expanded methylation data alone, both expression and expanded methylation data or both expression and the original 450K data on COAD (repeating for 100 times); (C) The hierarchical clustered heatmap using the selected features (both gene expression and methylated loci) in prognosis analysis; Both of the tumor samples and the features were clustered, and the log2 (RPKM) of gene expression value was normlized to [0,1]. (D) The Kaplan-Meier survival plot of the two clustered samples.

To validate whether the triple-evidenced genes could also perform well in other datasets. Firstly, we extracted gene expression data of normal and tumor tissues of liver, breast, uterus, bladder, esophageal and colon from GENT^30^, and tested the classification performances based on the triple-evidenced genes (Table 1); secondly, we extracted the expression data of 5 other cancers (STAD, READ, CHOL, GBM and PAAD) not included in the 13 cancers studied here from TCGA, and investigated the prediction performances based on the triple-evidenced genes (Table 2). We used the random forest model constructed from all the tumor samples with 47 features selected in more than half of the 100 cross validations. The prediction results showed satisfactory results and suggested that the triple-evidenced genes are important and robust for pan-cancer analysis.

**Table 1:**
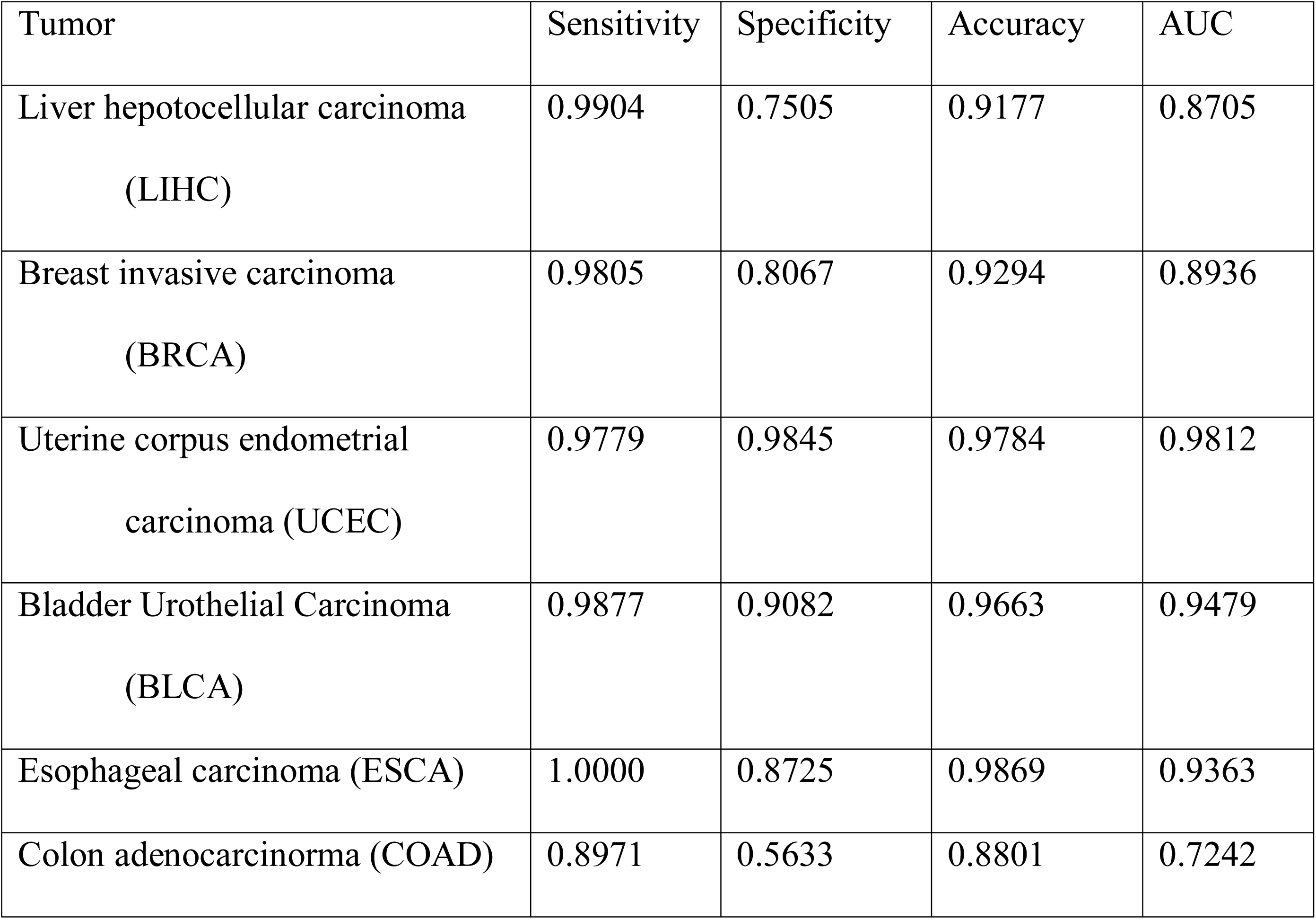
the classification performances on cancers from the GENT data

**Table 2:** the classification performances on 5 other cancers from the TCGA data

| Tumor | Sensitivity | Specificity | Accuracy | AUC |
| --- | --- | --- | --- | --- |
| Stomach Adenocarcinoma<br>(STAD) | 0.7734 | 0.8378 | 0.7791 | 0.8056 |
| Rectum adenocarcinoma<br>(READ) | 0.8842 | 1.0000 | 0.8942 | 0.9421 |
| Cholangiocarcinoma (CHOL) | 0.9722 | 1.0000 | 0.9778 | 0.9861 |
| Glioblastoma Multiforme<br>(GBM) | 0.9341 | 0.6211 | 0.9244 | 0.7671 |
| Pancreatic adenocarcinoma<br>(PAAD) | 0.8245 | 0.5278 | 0.8042 | 0.7122 |

Furthermore, we investigated whether it is possible to distinguish individual cancers. The candidate features were the combination of all the features selected in any of the 100 cross validations in the above diagnosis analysis. For the 13 cancer samples, multi-class logistic regression model was constructed based on the gene expression and the methylation levels of promoter CpGs using LASSO. The average prediction accuracy with 10-fold cross validation for 100 times was 95.27%±0.64%. This accurate prediction indicates that the expression and methylation features based on the triple-evidenced genes reflect the differential patterns not only between cancer and normal samples but also between different cancers. Among the 13 cancers, THCA and PRAD were with the highest accuracies (99.32% and 99.16%), while the LUSC was with the lowest accuracy (87.41%). When looking into the misclassification results, KIRP is prone to be predicted as KIRC and vice versa, LUAD is prone to be predicted as LUSC and vice versa, which is reasonable because they belong to the same tumor category. Also we found that the majority of mis-classified HNSC were predicted as LUSC, and many of the mis-classified LUSC were predicted as HNSC. The interesting results were consistent with the previous reports that patients treated for head and neck squamous cell carcinoma frequently developed second primary tumors in the lung, and they shared many common patterns^31, 32^.

Among the top 20 features (Table S5), 13 features are gene expression and 7 are CpG methylation values of the triple-evidenced genes. Some of these features have literature evidences to support their importance in discriminating cancers. For example, the gene expression of SUSD2 is the second most important feature, which is consistent with its reported variable expression in cancers, e.g. down-regulation in colon cancer^33^ and hepatocellular carcinoma^34^, and highly expressed in breast cancer^35^. Another example is CYGB expression, the fifth most important feature. CYGB shows variable expression in cancers: it is down-regulated in many cancers ^36^ but up-regulated in lung and brain metastases, and head and neck cancer ^37, 38^. The methylation level at a LAMA4 promoter CpG was found as the seventh most important feature; previously, the aberrant methylation at the LAMA4 promoter was observed in breast carcinoma^39^ and low methylation was associated with poor progression-free survival^40^.

### Prognostic value of the triple-evidenced genes

We also investigated whether the triple-evidenced genes are useful in predicting survival rate. For the survival data of each cancer (11 cancers with sufficient samples were analyzed, see details in Materials and Methods), we applied the LASSO cox proportional hazards regression for feature selection. The candidate features include gene expression and expanded methylation data (the methylation level of all the CpGs in the promoters) of triple-evidenced genes identified in each cancer. The performance was assessed using 10-fold cross validation for 100 times. We compared the concordances (C-indexes) based on four different candidate features (expanded methylation and expression levels, only expanded methylation, only expression level, and the original 450K methylation and expression levels of the triple-evidenced genes). As an example, the boxplots of the concordances of COAD in cross validation are shown in Figure 4B. The concordance based on the features selected from using both expression and expanded methylation data is superior to using either data alone, or the combination of the original 450K methylation and expression levels: the p-values are 0.03 (compared with expanded methylation data), 1.2e-11 (compared with expression data) and 9.1e-6 (compared with the combination of the original 450K methylation and expression levels).

Furthermore, we focused on the gene features frequently selected among the cross validations to cluster the tumor samples. For example, using the 19 features selected in more than 20% of the cross validations in the COAD samples (as shown in Figure 4C), hierarchical clustering identified two obvious subgroups, which shows significantly different survival times in the Kaplan-Meier survival plot in Figure 4D (p-value=0.00145). The results for the other 10 cancers are shown in Figure S10-S19. In 8 out of the 11 cancers, the concordances based on the features selected from the combination of both expression levels and expanded methylation data are the highest, indicating the usefulness of the expanded methylation data in prognosis analysis. Among the most often selected features, TNXB, RRM2, CELSR3, DBNDD1 and SLC16A3 are the top 5 most often selected genes in the survival analysis among the 11 cancers (Table S6 and see discussion below).

### Triple-evidenced genes important for both diagnosis and prognosis

Nine genes are most often selected in both diagnosis and prognosis analyses: TNXB, RRM2, CELSR3, SLC16A3, FANCI, MMP9, MMP11, SIK1, TRIM59. Their differential expression and methylation levels between normal and cancer samples as well as the somatic mutations in the 13 cancers are shown in Figure 5. The expression and somatic mutation patterns of the 9 genes are quite consistent in the 13 cancers but the methylation patterns of their promoters vary. These 9 genes are currently considered as biomarkers or potential biomarkers for diagnosis or prognosis in specific cancers. However, our analyses suggested that they are likely general biomarkers for at least the 13 cancers analyzed here.

**Figure 5.**
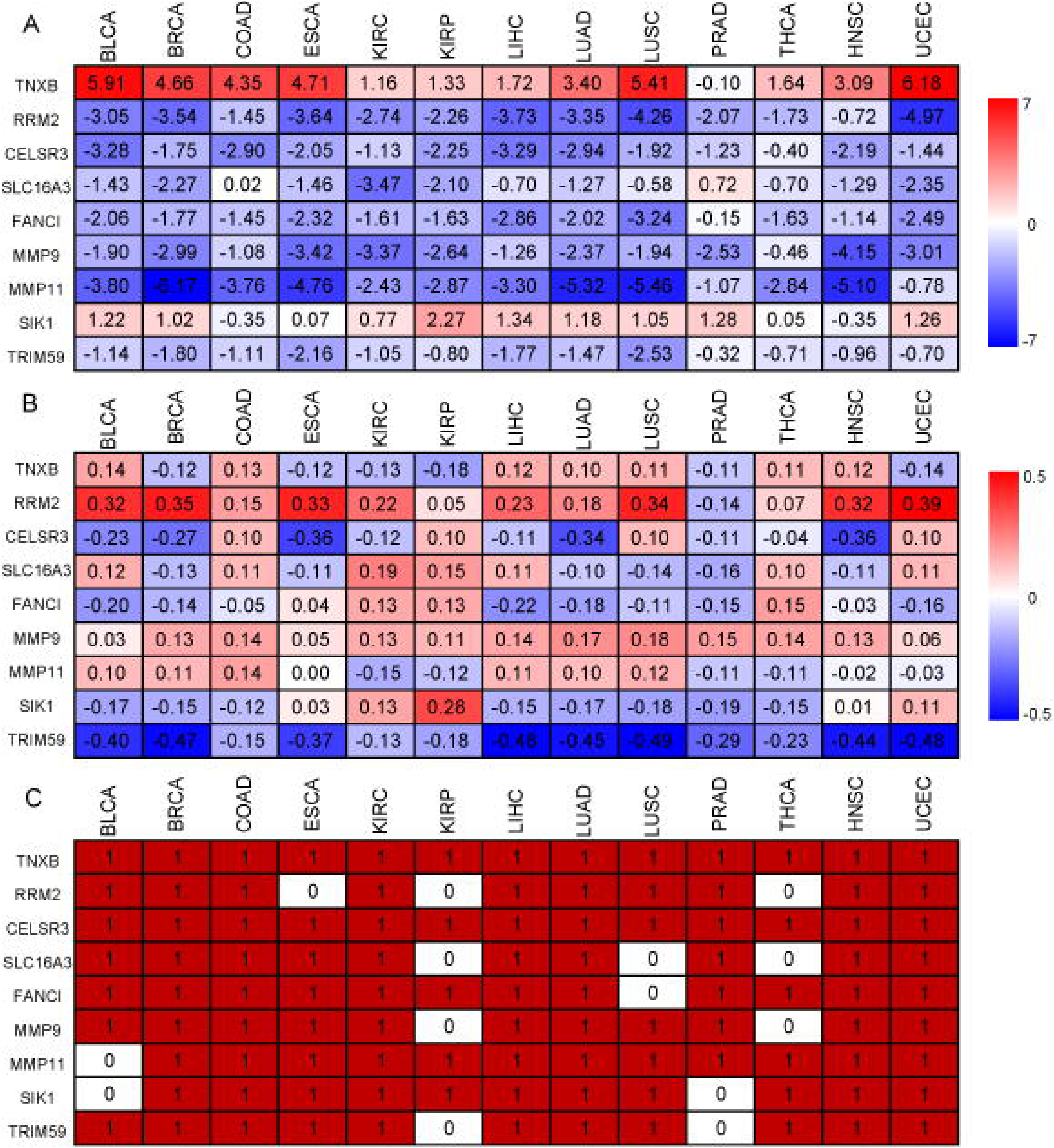
9 genes important for both diagnosis and prognosis analyses in the 13 cancers. (A) gene expression, (B) methylation and (C) somatic mutations of the 9 genes.

TNXB was reported as a potential marker for prognosis in patients with stage III serous ovarian cancer ^41^. RRM2 was reported as independent negative prognostic marker for survival in patients with resected pancreas cancer^42^ and a promising prognostic biomarker and therapeutic target for ER-negative breast cancer patients^43^. CELSR3 was suggested as a biomarker in OSCC prognostication^10^, and prognostic marker in small intestinal neuroendocrine tumor^5^. MMP11 and MMP9 were reported as breast tumor biomarkers and associated with tumor survival^28, 29^. For SLC16A3, there is no report on its prognostic power but studies showed that it might be an epigenetic marker for clinical outcome in clear cell renal cell carcinoma^44^.

It is worth of noting that FANCI and SIK1 genes could not be identified as the triple-evidenced genes using the original 450K array data. FANCI belongs to Fanconi anemia complementation group and it was a negative regulator of Akt activation that connects with the oncogenic PI3K-Akt pathway and the tumor suppressing FA pathway^45^. This gene has also been linked to drug resistance in cancer treatment^46^. SIK1 is stimulated by a cancer suppressor LKB1, which leads to metastatic spread and invasiveness, as well as apoptosis resistance^47^. Loss of SIK1 has been found in epithelial ovarian cancer and pancreatic cancer^48^, and decreased SIK1 expression is correlated with poor outcome of breast cancer treatment^49^, indicating the potential application in prognosis. Our results further support the potential of using FACNI and SIK1 as prognosis marker and provide insight in broadening its application in other cancer types.

Among the 9 genes, RRM2, MMP9, MMP11 and SIK1 are known drug targets. For example, they are inhibited by gemcitabine (RRM2), marimastat (MMP9 and MMP11) for treating several cancers. Our analyses suggested these inhibitors may be effective for the majority of the 13 cancers, which suggests possible broader applications of these inhibitors. For example, gemcitabine targeting RRM2 are validated for the treatment of non small cell lung cancer^50^, ovarian cancer^51^, pancreatic cancer^52^, adrenocortical cancer^53^, and oral squamous cell carcinoma^54^. We speculate that gemcitabine can be used to treat bladder, colon, kidney, liver and prostate cancers. Furthermore, HG-9-91-01(SIK1) was reported to induce anti-inflammatory phenotype and could be used to treat certain autoimmune diseases ^55, 56^, we speculate that it can be repurposed to treat cancers as 4 of the top 5 enriched pathways in the pan-cancers analysis are closely related to inflammatory processing or inflammation response.

## Discussion

We present here a method EAGLING to significantly expand the Illumina 450K array data with a fast speed and better precision than the previous models. We have performed pan-cancer analysis on 13 TCGA cancers to identify genes with differential methylation and gene expression between cancer and normal samples as well as containing somatic mutations. These triple-evidenced genes, particularly TNXB, RRM2, CELSR3, SLC16A3, FANCI, MMP9, MMP11, SIK1 and TRIM59 show diagnostic and prognostic power. Note that FANCI and SIK1 could only be identified as triple-evidenced genes using the expanded methylation data. The pathways in which they are enriched also suggest new therapeutic targets or repurposing the existing drugs. We focused on discussing the common features among the 13 cancers but it is worth of noting that the triple-evidenced genes in individual cancers can also be potential biomarkers or drug targets.

## Conclusion

We showed that the expanded methylation data allowed identification of more cancer-related genes, which led to better performance in both diagnosis and prognosis. The common patterns shared among the 13 cancers suggest that some drugs (such as gemcitabine) currently aiming to specific cancers might be useful to treat other cancers, and drugs aiming to immune diseases (such as HG-9-91-01) might be repurposed for cancer therapy.

## Methods

### DNA methylation data for model construction

In total, 33 tissues or cell lines with both WGBS and 450K array data were retrieved from the NIH Roadmap Epigenomics project^57^ and TCGA(Table S1). We downloaded the methylation proportion values of WGBS data and beta values of 450K array from the GEO Database and TCGA data portal directly. Both WGBS data and 450K array data were quantile normalized. We used the quantile normalization between the 450K arrays, and between the WGBS data, to reduce the batch effect, as the quantile normalization strategy was reported to be efficient for the intra- and inter-arrays normalization^58, 59^.

In the EAGLING model, the CpG site to be predicted is denoted as L. The WGBS methylation value at L in the tissue that shows the most similar local methylation profile among the 33 tissues was used as one feature (*x*_1_) and the methylation value measured by the Illumina 450K array of the closest neighbor CpG of L was used as the second feature (*x*_2_). The local methylation profile was defined by 4 CpGs in the upstream and downstream 30kbp regions of site L (see Results and Figure 1 for details of how these parameters were selected). Only the CpG loci having 4 neighbor CpGs in the 30kbp flanking regions would be considered for expansion. A logistic regression model was built on these two features to predict the methylation level at L. The main differences between EAGLING and our previous model ^3^ include: (1) not using DNA sequence features to achieve a faster speed, (2) optimized parameters of local methylation pattern and the flanking region size, (3) the training set was significantly increased from 14 to 33 that is expected to improve the performance.

The performance of EAGLING was assessed by leave-one-tissue-out cross validation on all the 22 autosomes. The evaluation metrics included Pearson correlation coefficient (COR), Concord (CONCORD, the percent of CpGs with a methylation proportion difference less than 0.25 ^60^), Sensitivity (SE), Specificity (SP), Accuracy (ACC) and AUC (Area Under ROC Curve). For calculating SE, SP, ACC and AUC, we defined the methylation status as +1 if the methylation value is larger than 0.5, and the methylation status as -1 otherwise.

For performance validation, the 450K array and WGBS data of K562 and HepG2, two independent cancer cell lines from ENCODE project were retrieved from GEO database. The expanded methylation levels from EAGLING model were compared with the real WGBS data for performance validation.

### Expanding the DNA methylation data in the TCGA samples

We downloaded the 450K array data from TCGA. We only included 13 cancers with at least 10 normal samples of 450K array and RNA-seq data for the integrative analysis (Table S7): Lung adenocarcinoma (LUAD), Lung squamous cell carcinoma(LUSC), Breast invasive carcinoma (BRCA), Bladder Urothelial Carcinoma (BLCA), Colon adenocarcinorma (COAD), Kidney renal clear cell carcinoma (KIRC), Kidney renal papillary cell carcinoma (KIRP), Prostate adenocarcinoma (PRAD), Esophageal carcinoma (ESCA), Liver hepotocellular carcinoma (LIHC), Thyroid carcinoma (THCA), Uterine corpus endometrial carcinoma (UCEC), Head and neck squamous cell carcinoma (HNSC). All the 450K array data were quantile normalized.

In calculating the ratio of hyper/hypo-methylated CpGs of the tumor and normal samples, the methylation value(ranging from 0(totally unmethylated) to 1(totally methylated)) larger than 0.7 was defined as hyper-methylation, and the value less than 0.3 was defined as hypo-methylation. For comparison, the ratios of hyper/hypo-methylated CpGs of WGBS data of lung cancers were calculated.

### Identification of triple-evidenced genes in cancers

We compared the methylation levels of each CpG between tumor samples and the corresponding normal sample data, and defined a CpG site to be differentially methylated (DML) if the q-value of t-test <0.05 and the absolute difference of methylation value > 0.1. We considered genes whose promoters contain any CpG covered by the original or the expanded methylation data. To identify the genes differentially methylated in each cancer, the methylation status of all the CpG sites covered in promoters were considered. For each promoter, the Fisher’s combined test was used to get the q-value to evaluate whether a gene is differentially methylated. Similar to call DMLs, genes with q-value < 0.05 and mean difference of DNA methylation > 0.1 were selected as differentially methylated genes (DMGs).

The RNA-seq data were downloaded for the cancers listed in Table S7 from TCGA. The sample sizes are also shown in Table S7. The gene expression data were log2-transformed and normalized. Differentially expressed genes (DEGs) were defined if the fold change > 2 and the q-value of t-test is < 0.05.

To collect genes with mutation related to the 13 cancers, we downloaded the somatic mutation level 2 data from TCGA. For each cancer, a gene that was annotated with curated somatic mutation in TCGA was considered. We extracted the gene lists with somatic mutation of all the 13 cancers for integrative analysis.

### Diagnostic and prognostic analysis

To search for common features in pan-cancer analysis, all the 13 cancer samples and normal samples were combined together, respectively. A random forest model was constructed using cross validation. In each of the cross validation, the cancer and normal samples of 10 cancers were randomly selected for model training, and the remaining samples of 3 cancers were used for test. The cross validation was repeated for 100 times. We chose the triple-evidenced genes that appear in more than half of the 10 training cancer types as the candidate genes. Their associated gene expression and DNA methylation levels of individual CpGs in the promoters of the selected genes in the expanded data were used as input features; Both of the gene expression and the methylation levels of CpGs in the promoters were candidate features for feature selection with LASSO.

To indicate whether these potential common features also reflect some differences between the cancers, a multi-class regression model was constructed with the tumor samples of the 13 cancers. The candidate features were the combination of all the features selected in any of the 100 cross validations in the above diagnosis analysis. The gene expression and the methylation levels of promoter CpGs were further selected using LASSO.

In the prognostic analysis, the PRAD and THCA were not analyzed due to their limited samples with the expression, methylation and survival data. The sample sizes of the 11 tumors are listed in Table S8. For each of the remaining 11 tumors, both of the expression levels and DNA methylation levels of CpGs in the promoter regions were included for variable selection with LASSO Cox proportional hazards regression model, and the concordances in the 10-fold cross validations were compared for prognostic power.

## Data Availability

The datasets analysed during the current study are available in the TCGA (https://cancergenome.nih.gov/) and NCBI’s GEO (https://www.ncbi.nlm.nih.gov/geo/). The R scripts for the EAGLING model and the triple-evidenced genes are available at http://114.55.236.67:8013/Integrative_Analysis/home.

### Acknowledgements

This work was supported by the NIH(R01 HG009626), National Natural Science Foundation of China(61503061 and 61872063); and Fundamental Research Funds for the Central Universities(ZYGX2016J102); and the open fund of Key Laboratory of Symbolic Computation and Knowledge Engineering of Ministry of Education(93K172017K02).

## Contributions

SF and WW conceived the study. SF, JT, NL, YZ, RA, KZ, MW and WD performed the analysis. SF and WW wrote the manuscript with support from all authors. All authors read and approved the final manuscript.

