## Supplementary File for "Integrative analysis with expanded DNA methylation data reveals common key regulators and pathways in cancers"

**Supplementary Materials**

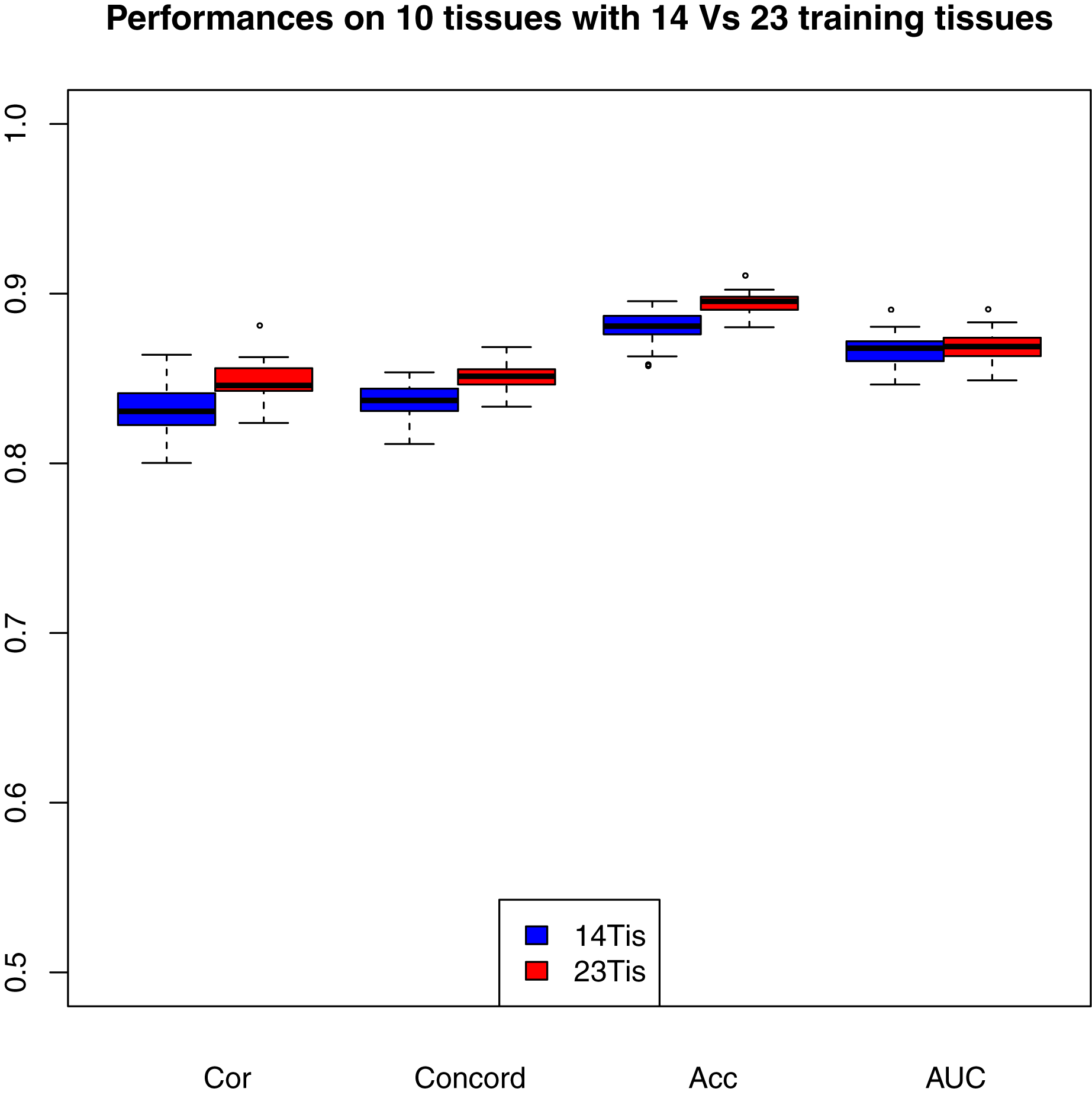

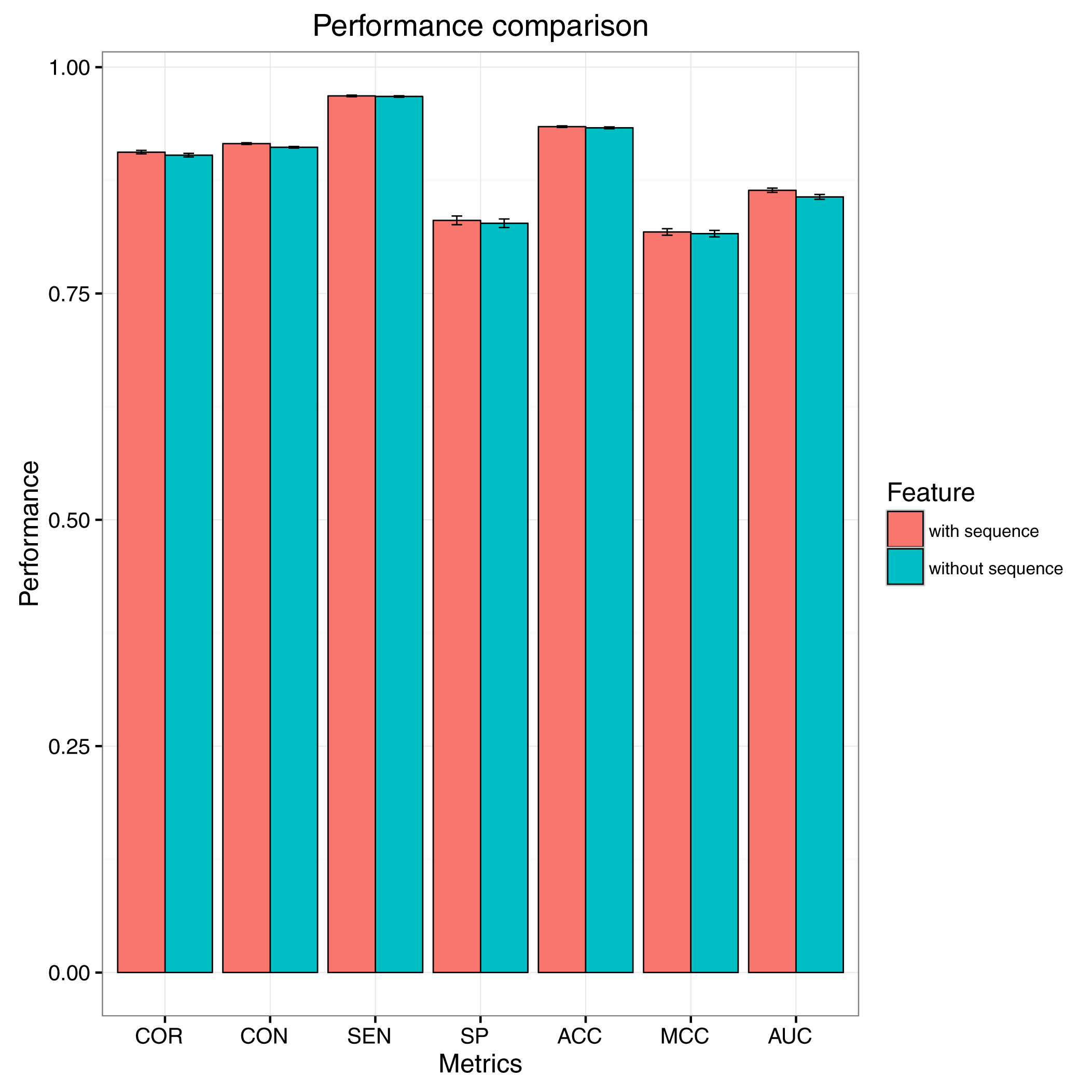

A	B

Figure S1. Comparison between EAGLING and the previous model [1]. (A). Inclusion of more training data improves the performance (repeating for 10 times); (B) Not including DNA sequence features does not affect the performance (Leave-one-tissue-out cross validation).

Figure S2. Differentially expressed genes (DEG), differentially methylated genes (DMGs) and genes containing somatic mutations in the 12 cancers. The 12 caners include BRCA: Breast invasive carcinoma; BLCA: Bladder Urothelial Carcinoma; COAD: Colon adenocarcinorma; ESCA: Esophageal carcinoma; HNSC: Head and neck squamous cell carcinoma; KIRC: Kidney renal clear cell carcinoma; KIRP: Kidney renal papillary cell carcinoma; LIHC: Liver hepotocellular carcinoma; LUAD: Lung adenocarcinoma; PRAD: Prostate adenocarcinoma; THCA: Thyroid carcinoma; UCEC: Uterine corpus endometrial carcinoma.

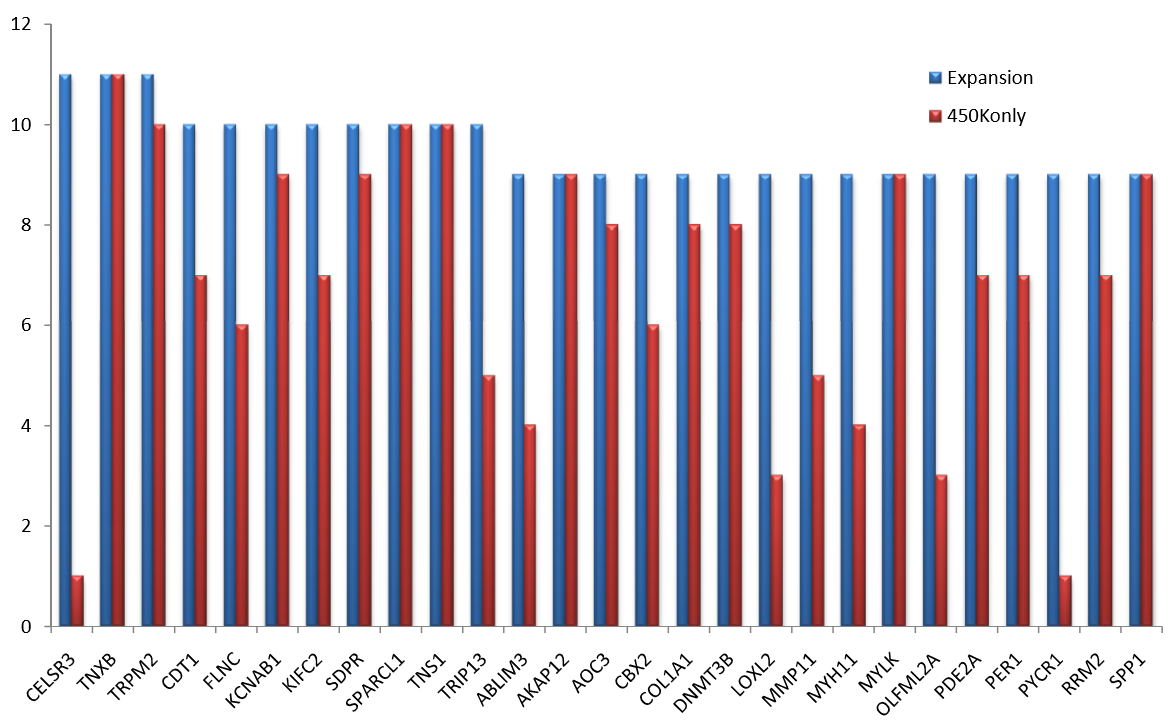

Figure S3. The important triple-evidenced genes could be identified in more cancers using the expanded methylation data compared to using the original 450K array data.

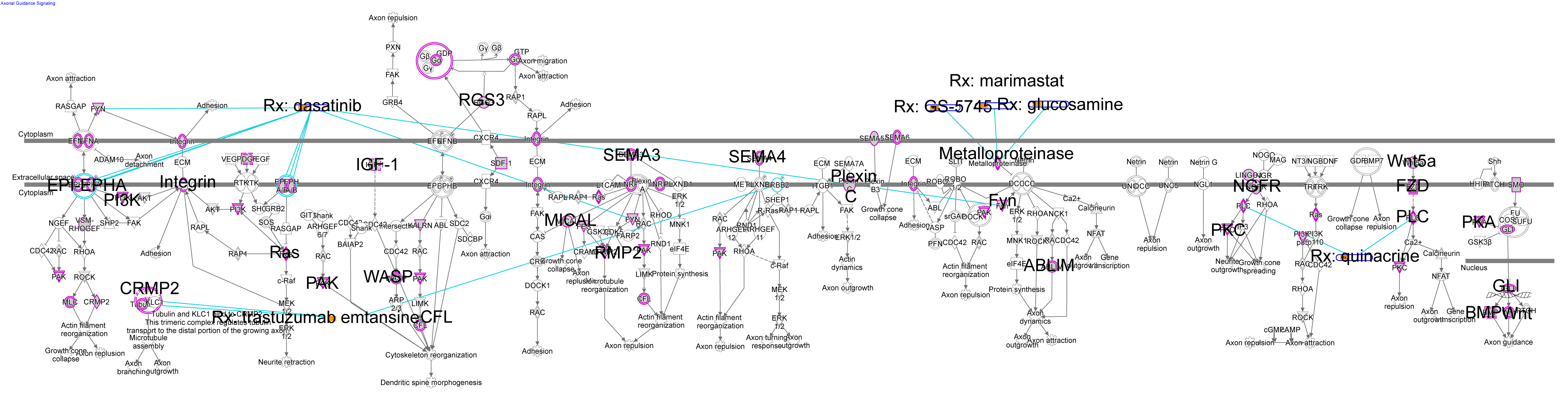

Figure S4. Axonal guidance signaling pathway appears to be important in many cancers. Triple-evidenced genes in LUSC are marked with purple circle; the top genes shared in the 11 cancers on the pathway are marked with star shape; some inhibitors or FDA approved drugs and their target genes are connected with blue lines.

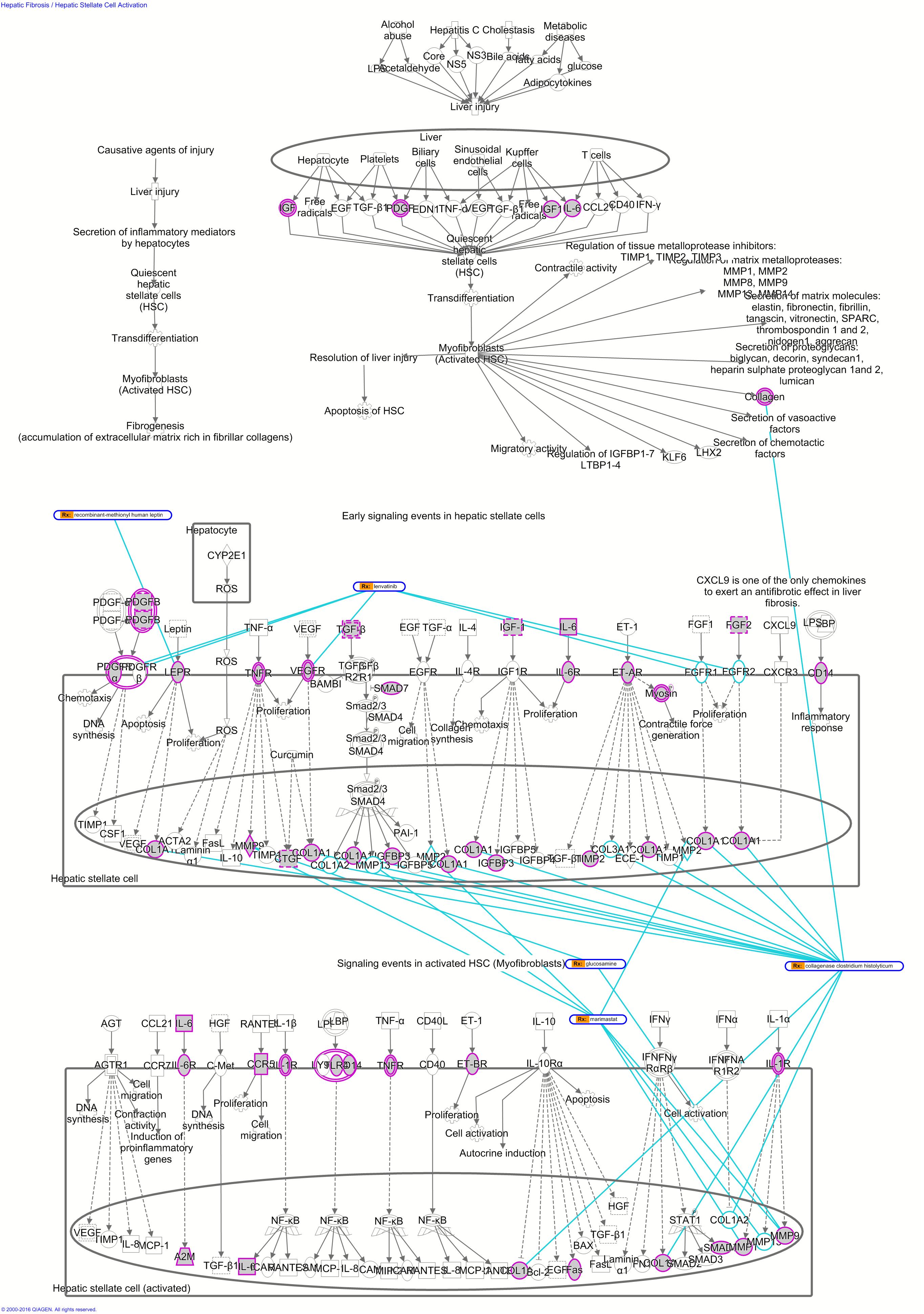

Figure S5. Hepatic Fibrosis/Hepatic Stellate Cell Activation pathways (LUSC). Triple-evidenced genes in LUSC are marked with purple circle; some inhibitors or FDA approved drugs and their target genes are connected with blue lines.

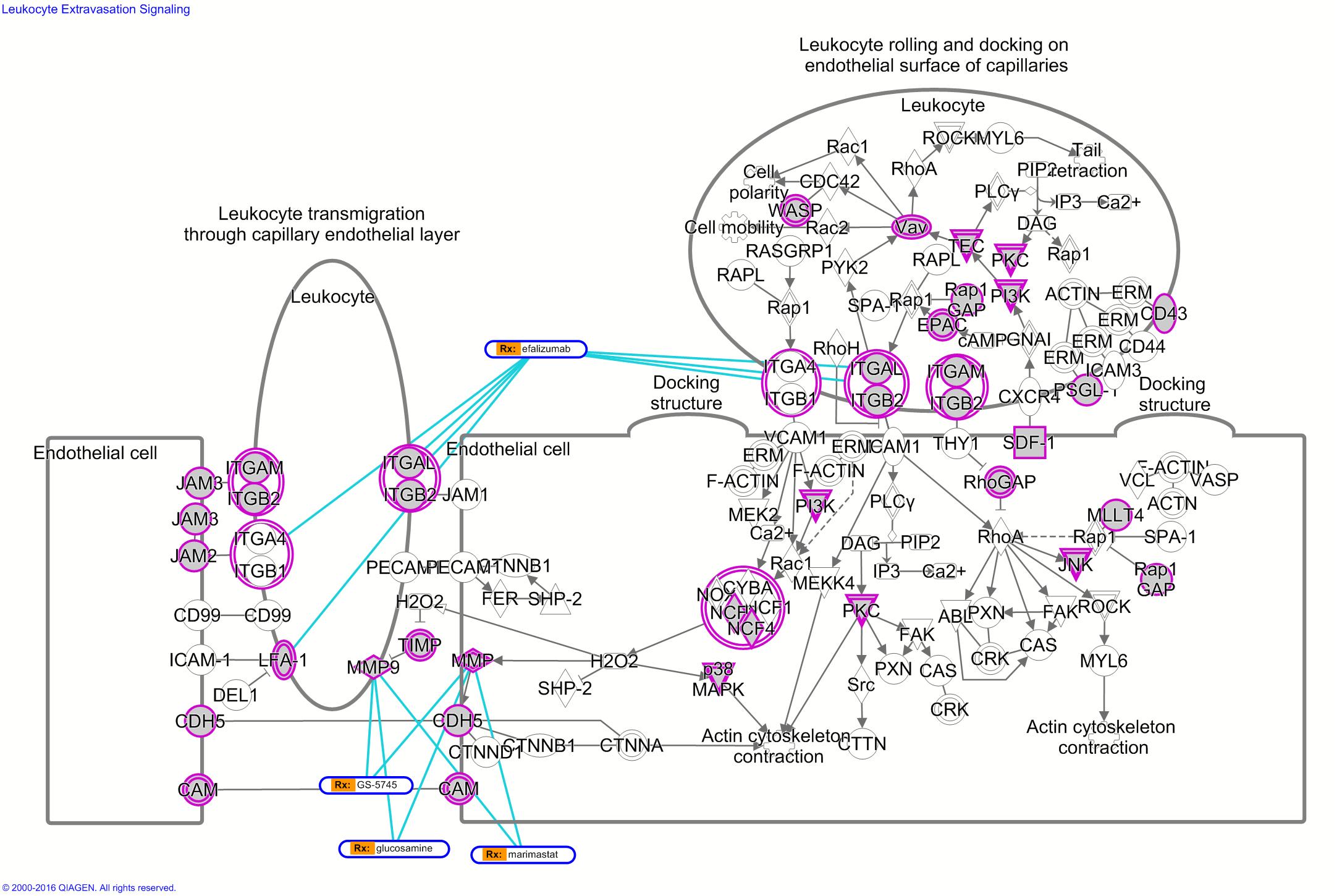

Figure S6. Leukocyte Extravasation Signaling (LUSC). Triple-evidenced genes in LUSC are marked with purple circle; some inhibitors or FDA approved drugs and their target genes are connected with blue lines.

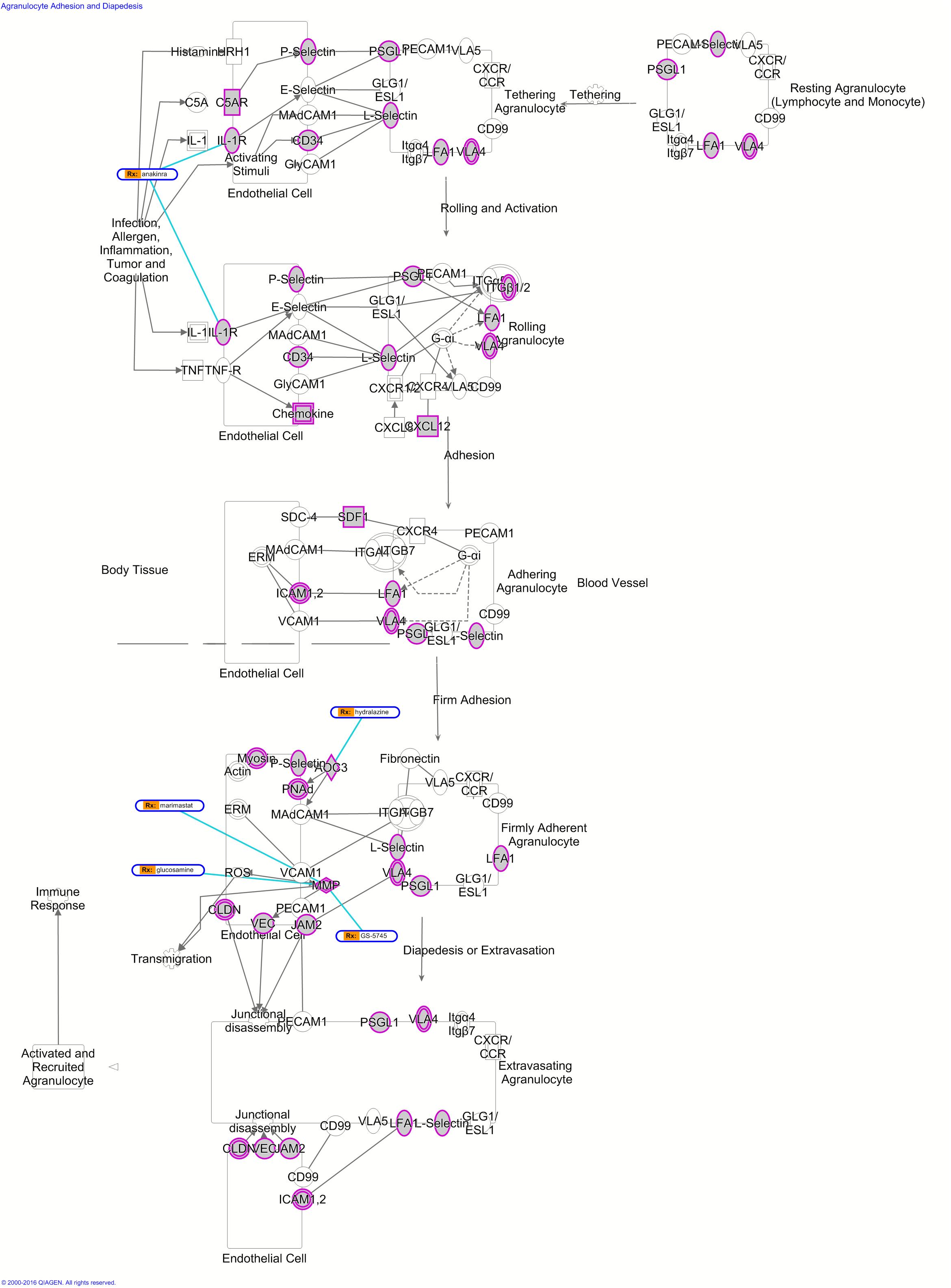

Figure S7. Agranulocyte Adhesion and Diapedesis (LUSC). Triple-evidenced genes in LUSC are marked with purple circle; some inhibitors or FDA approved drugs and their target genes are connected with blue lines.

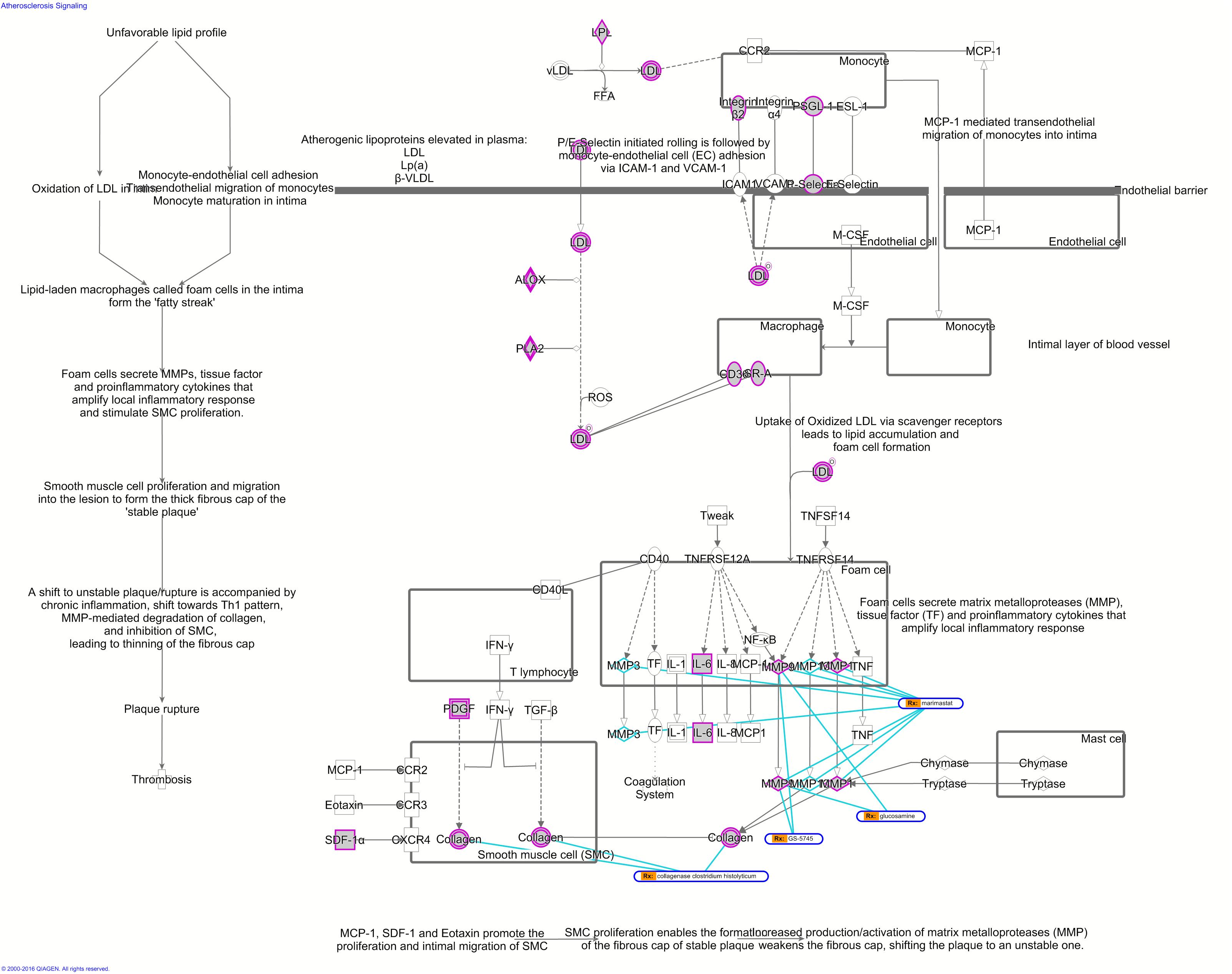

Figure S8. Atherosclerosis Signaling (LUSC). Triple-evidenced genes in LUSC are marked with purple circle; some inhibitors or FDA approved drugs and their target genes are connected with blue lines.

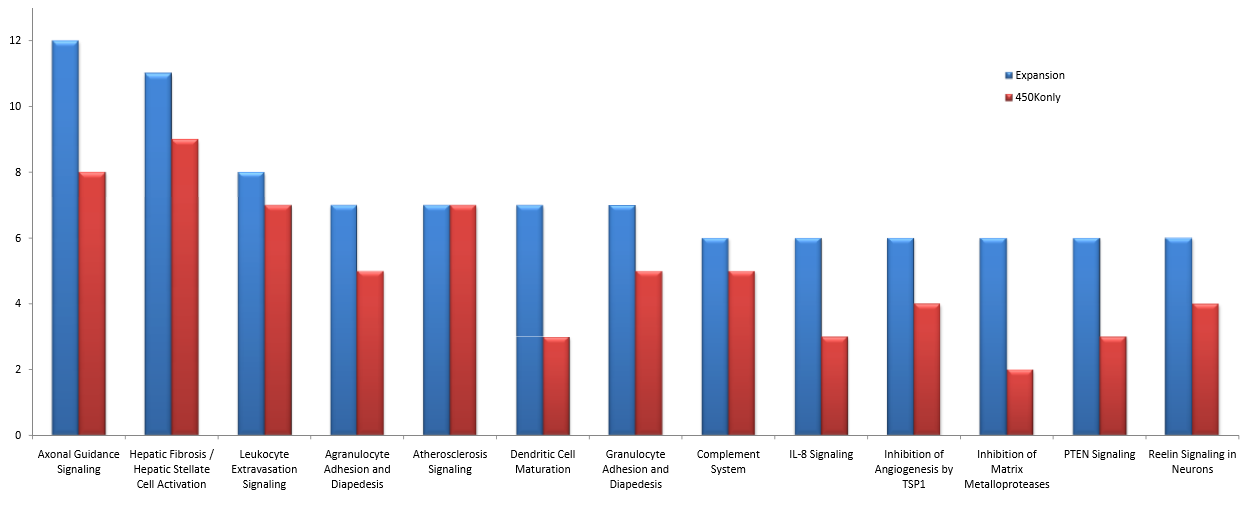

Figure S9. Important pathways could be identified in more cancers using the expanded methylation data than using the 450K array data.

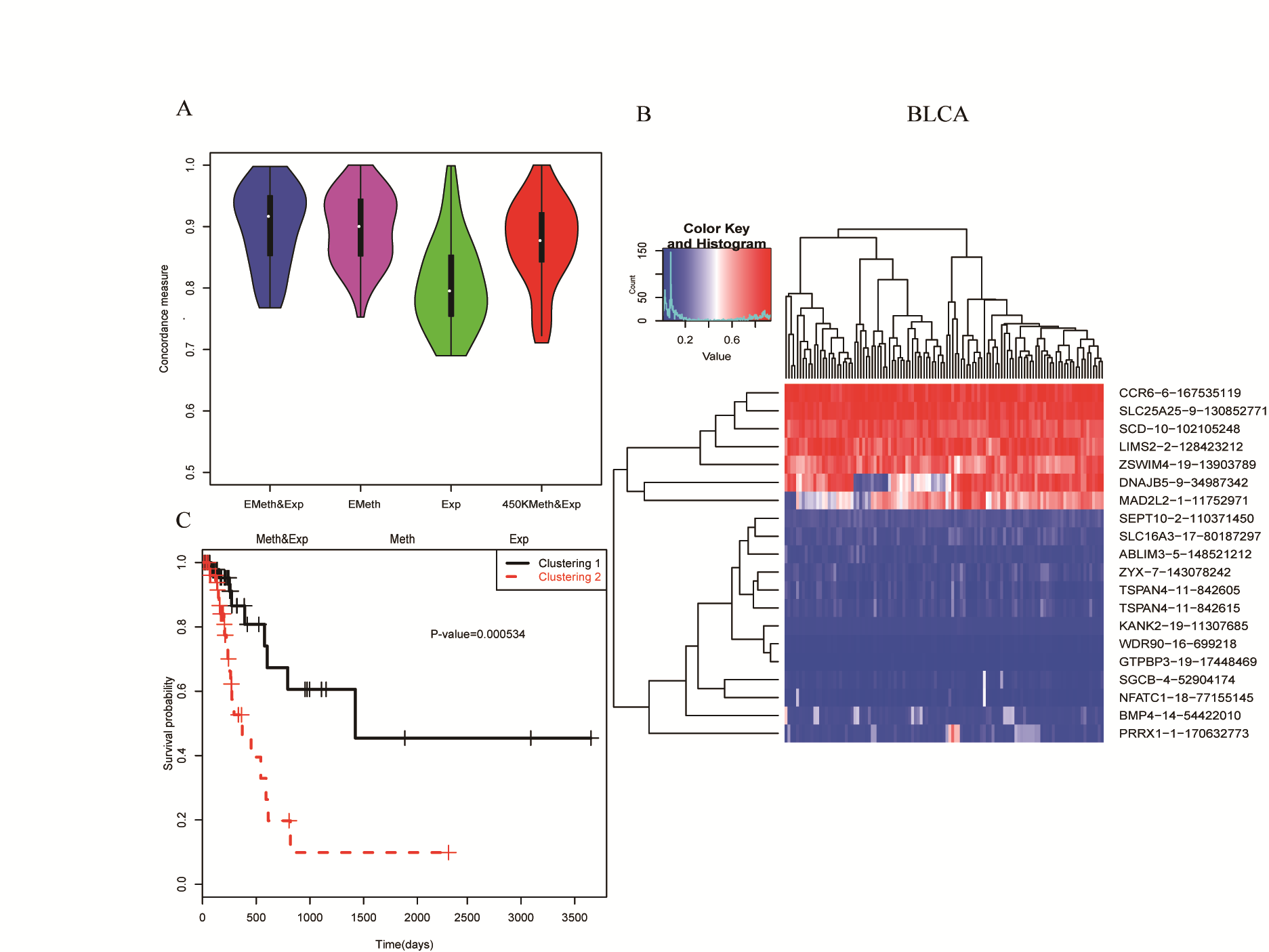

Figure S10. Survival analysis of BLCA using the features selected in more than 20% of the cross validations (repeating for 100 times). Each row is a selected feature (gene expression or methylation locus) and each column is a tumor sample.

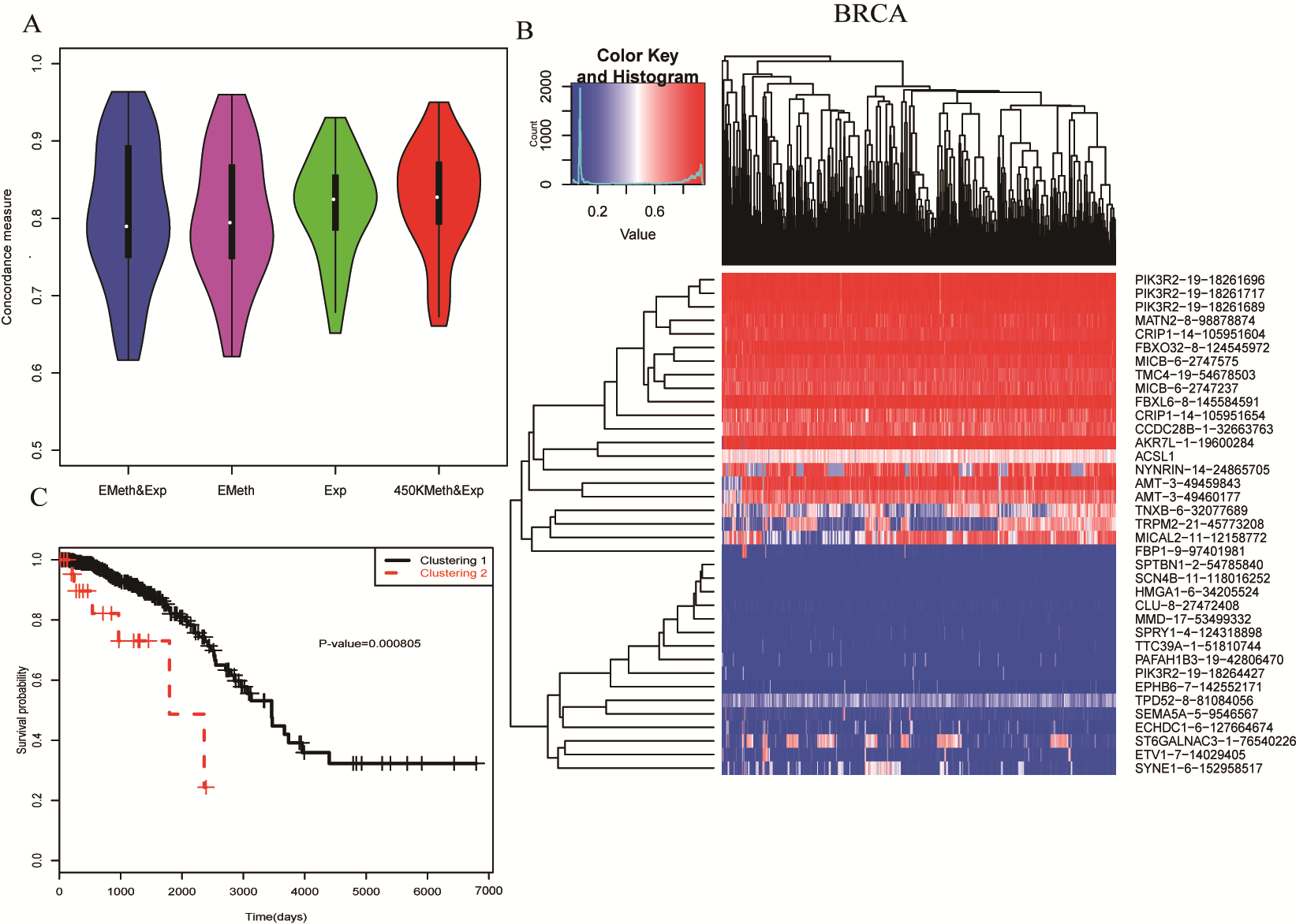

Figure S11. Survival analysis of BRCA using the features selected in more than 20% of the cross validations. Each row is a selected feature (gene expression or methylation locus) and each column is a tumor sample.

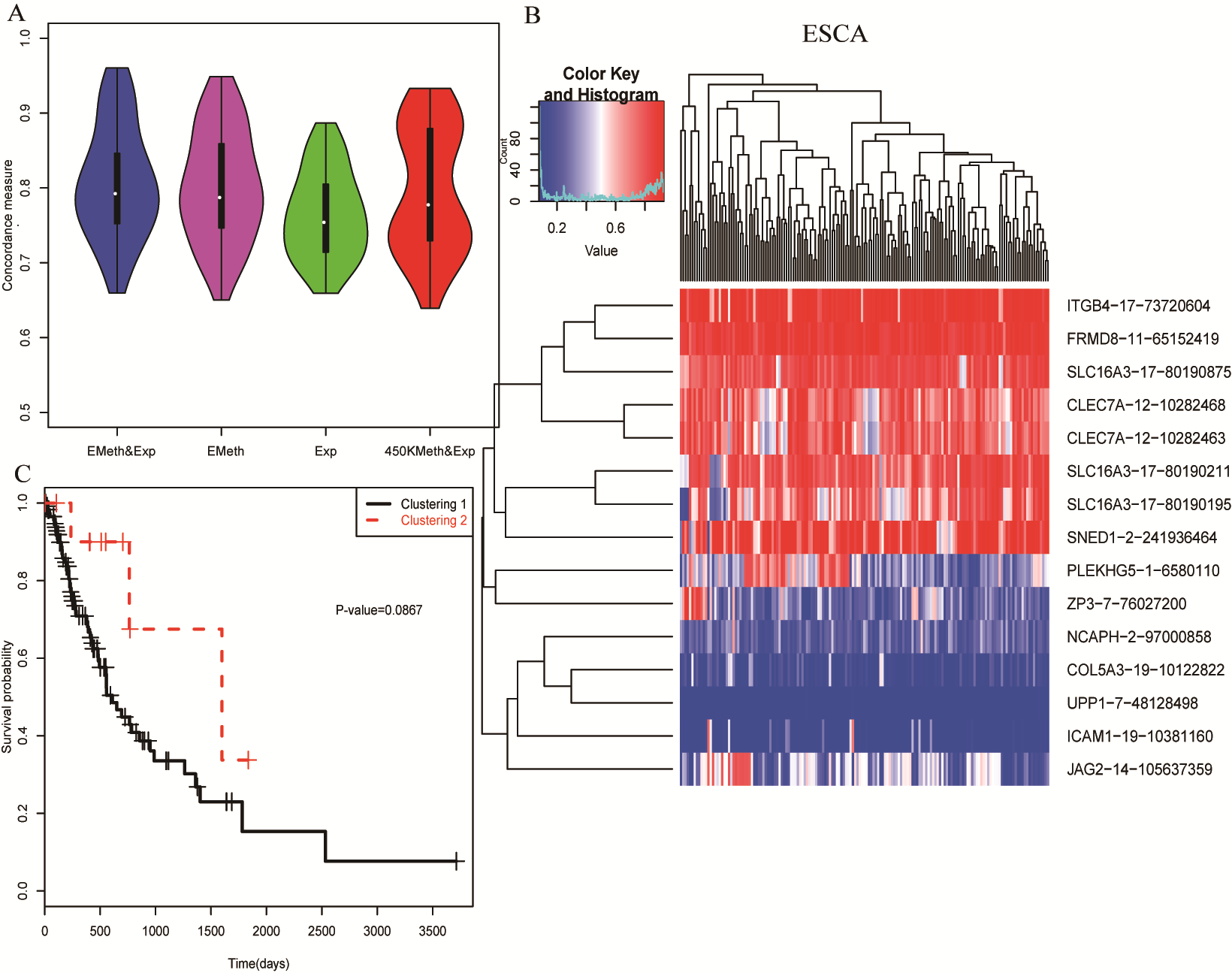

Figure S12. Survival analysis of ESCA using the features selected in more than 20% of the cross validations. Each row is a selected feature (gene expression or methylation locus) and each column is a tumor sample.

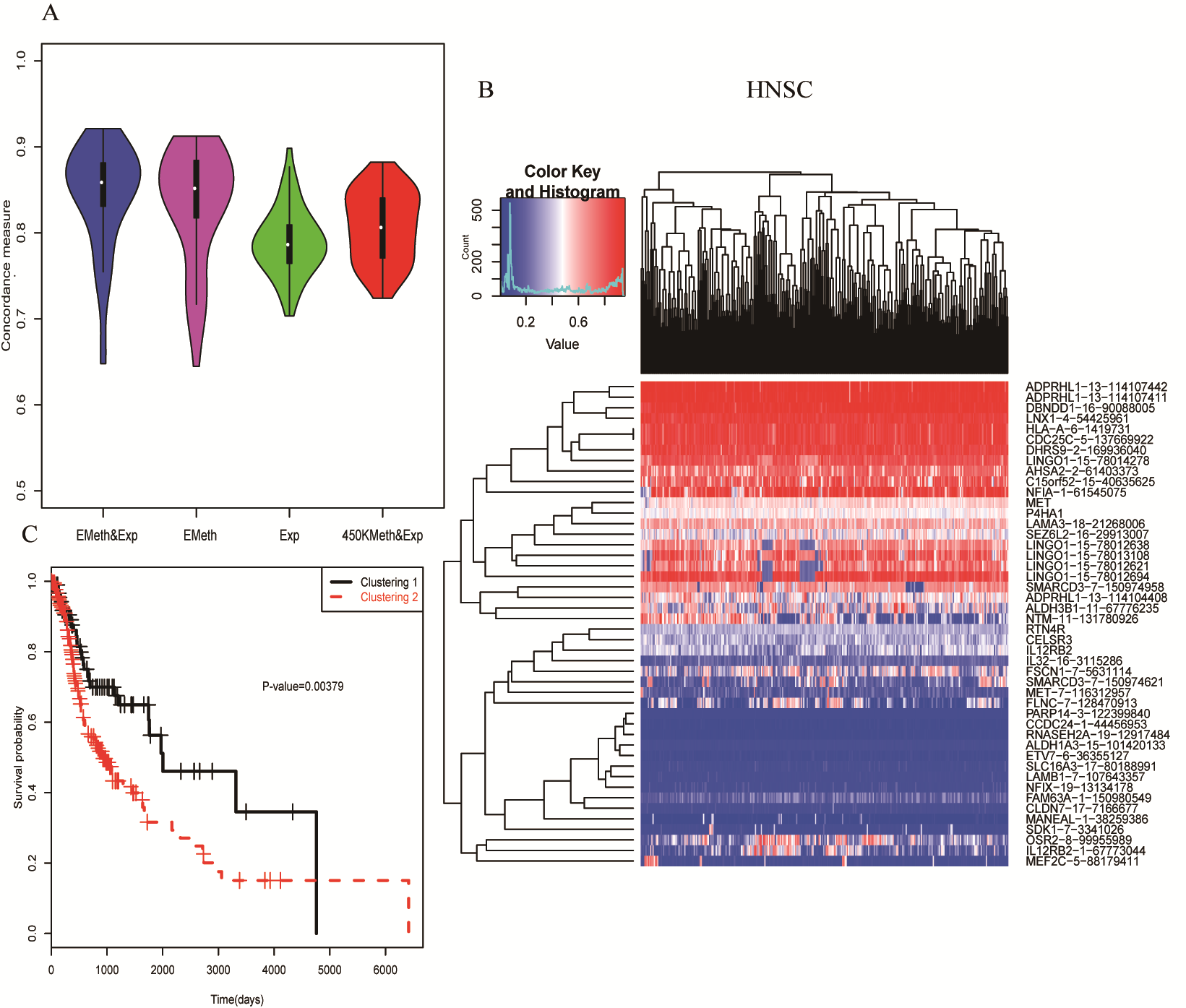

Figure S13. Survival analysis of HNSC using the features selected in more than 20% of the cross validations. Each row is a selected feature (gene expression or methylation locus) and each column is a tumor sample.

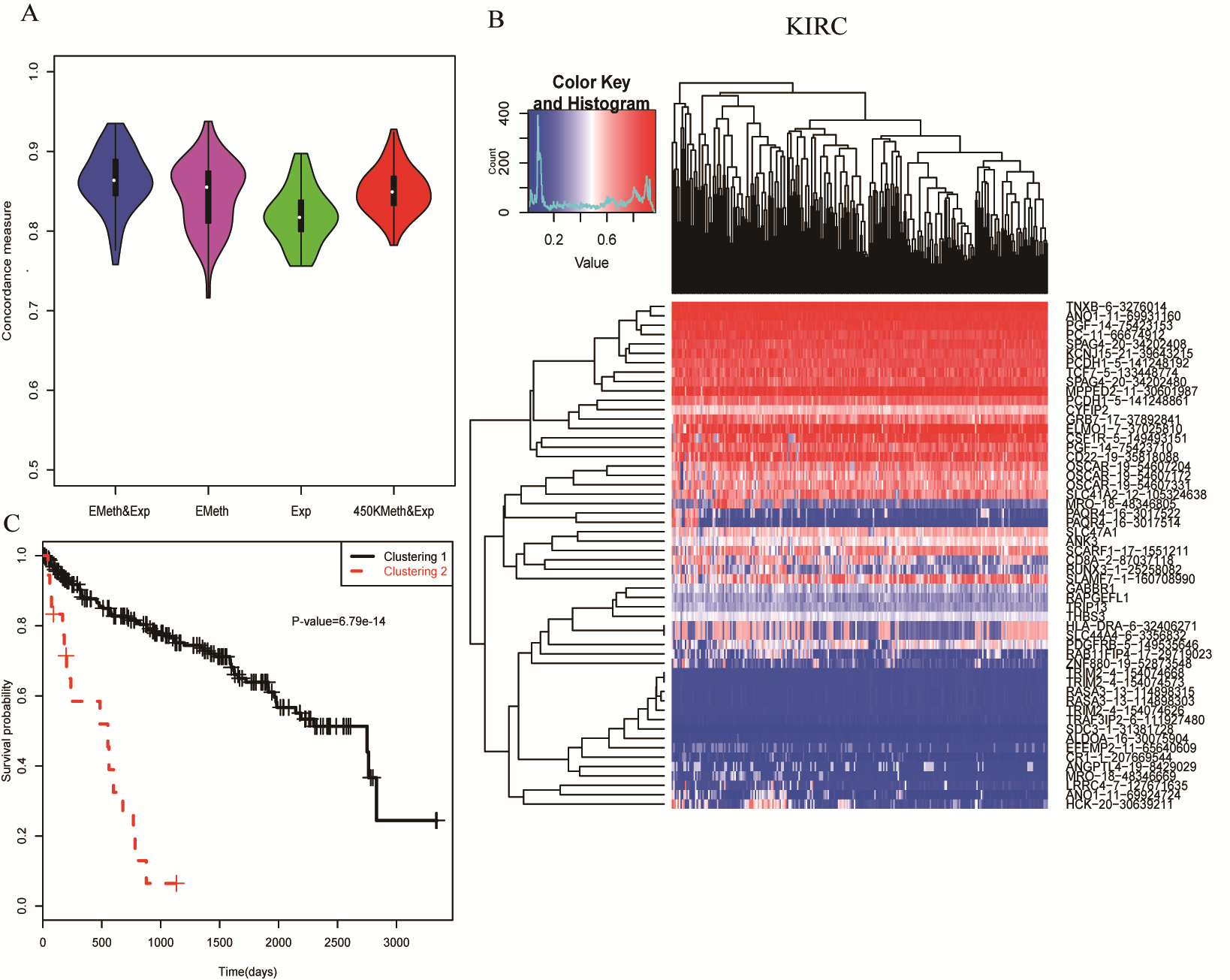

Figure S14. Survival analysis of KIRC using the features selected in more than 20% of the cross validations. Each row is a selected feature (gene expression or methylation locus) and each column is a tumor sample.

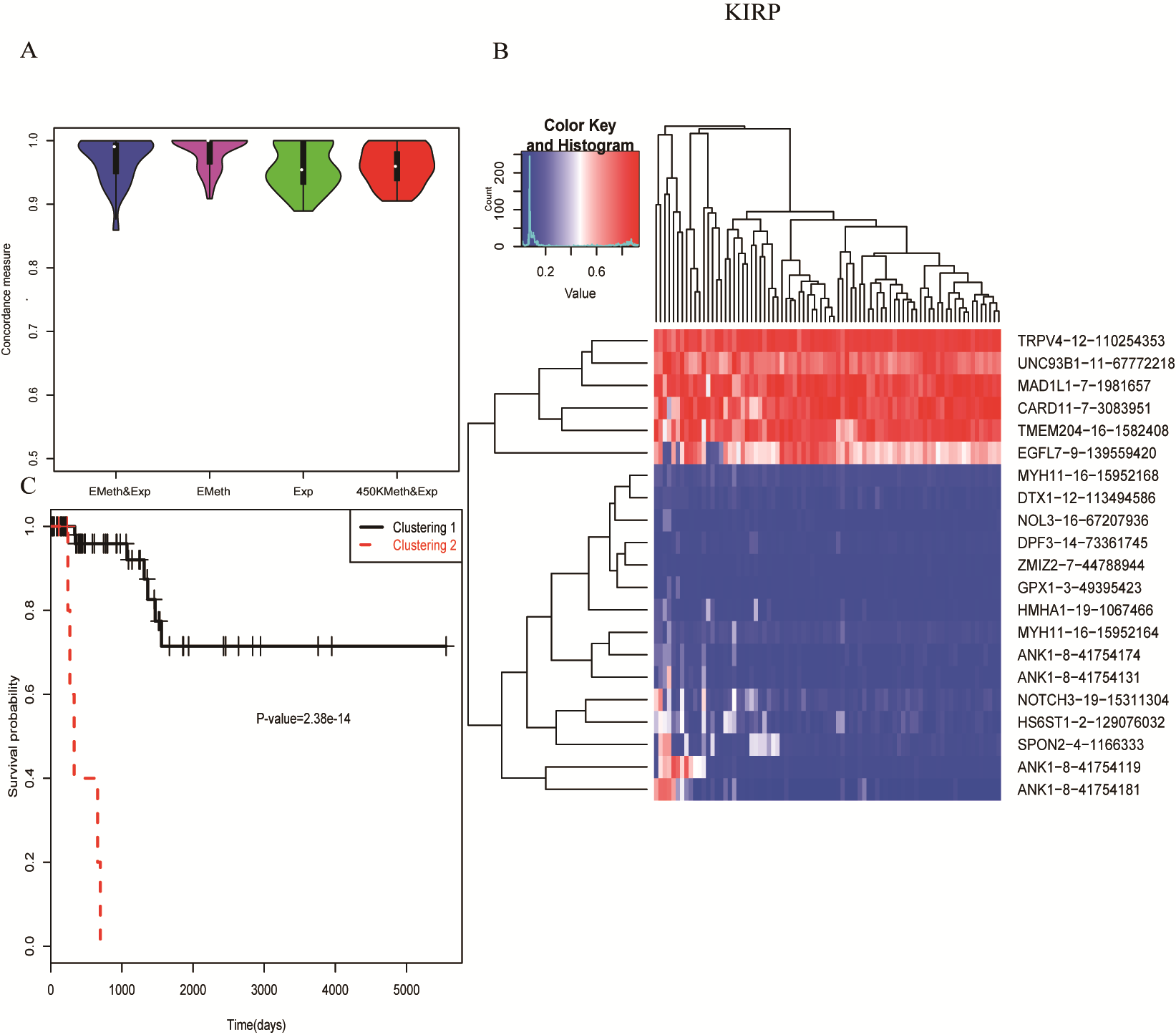

Figure S15. Survival analysis of KIRP using the features selected in more than 20% of the cross validations. Each row is a selected feature (gene expression or methylation locus) and each column is a tumor sample.

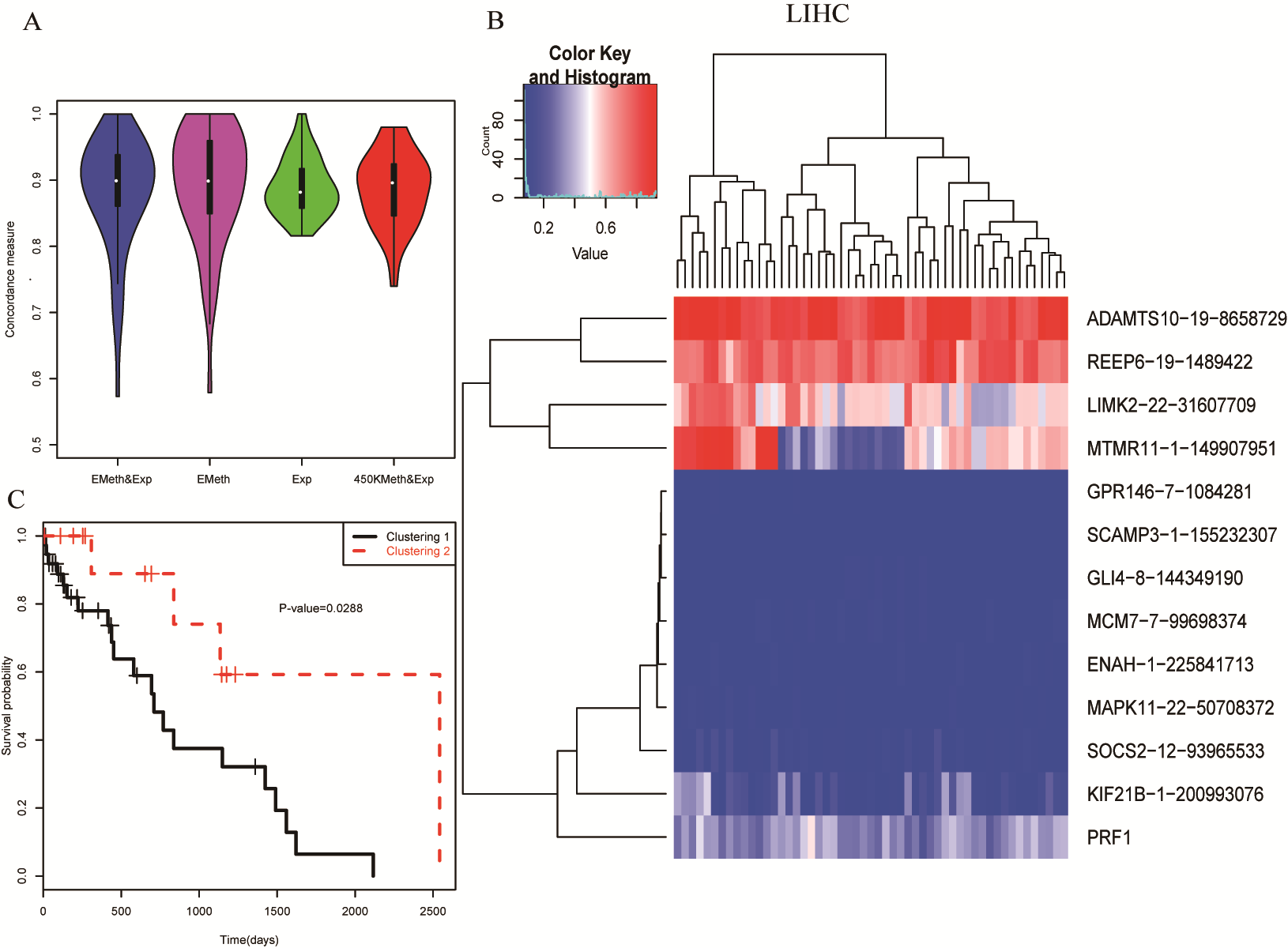

Figure S16. Survival analysis of LIHC using the features selected in more than 20% of the cross validations. Each row is a selected feature (gene expression or methylation locus) and each column is a tumor sample.

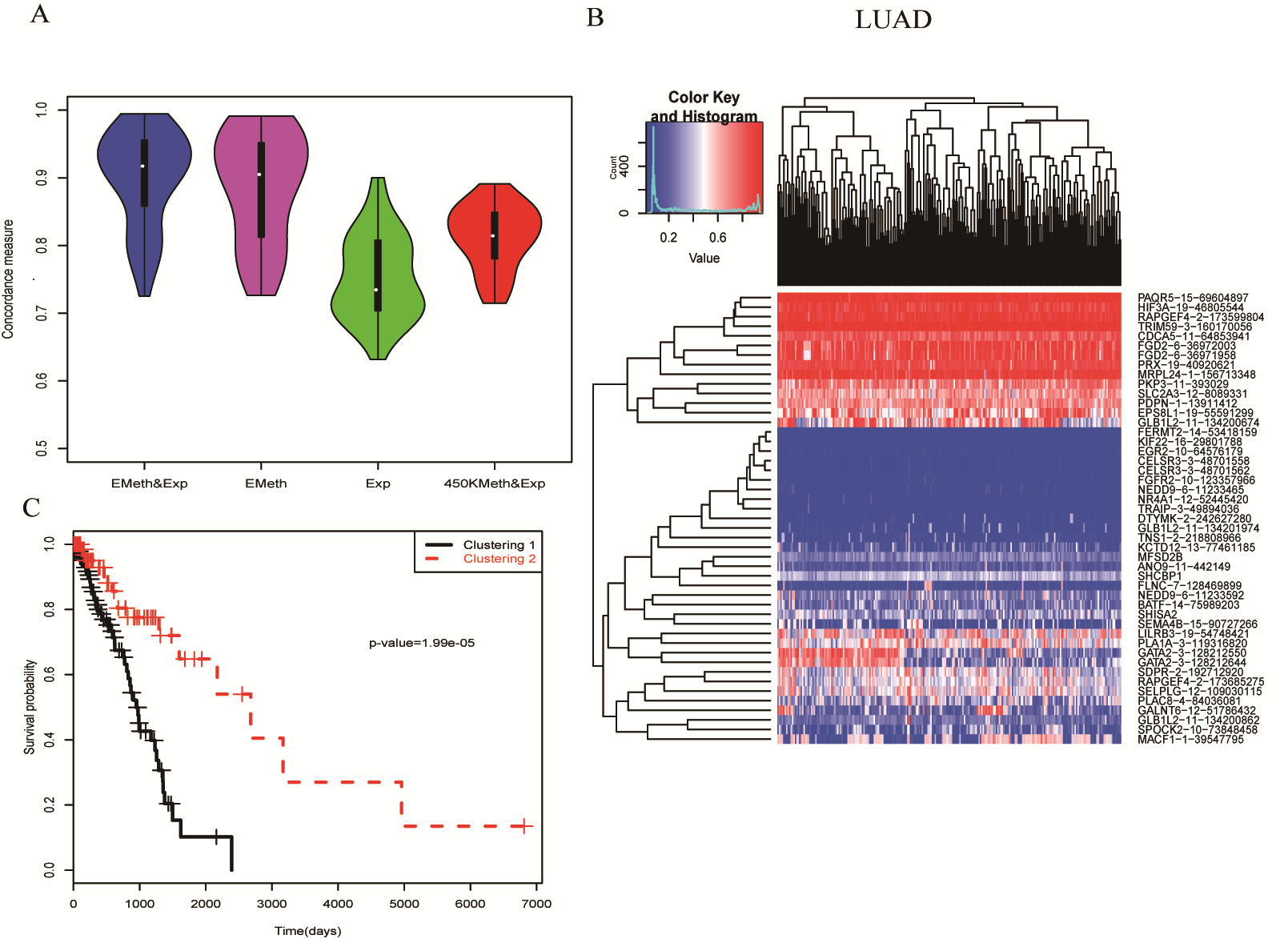

Figure S17. Survival analysis of LUAD using the features selected in more than 20% of the cross validations. Each row is a selected feature (gene expression or methylation locus) and each column is a tumor sample.

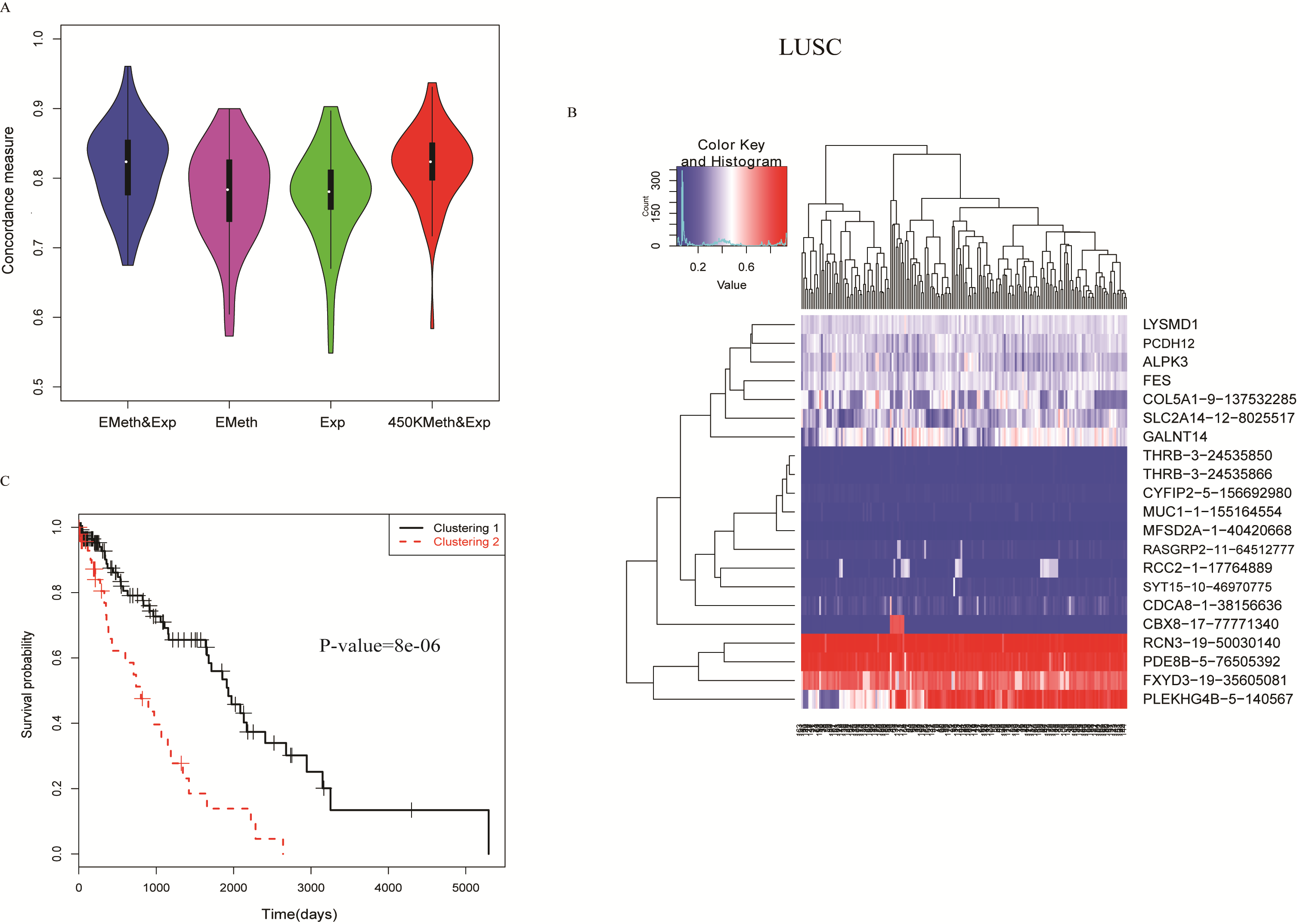

Figure S18. Survival analysis of LUSC using the features selected in more than 20% of the cross validations. Each row is a selected feature (gene expression or methylation locus) and each column is a tumor sample.

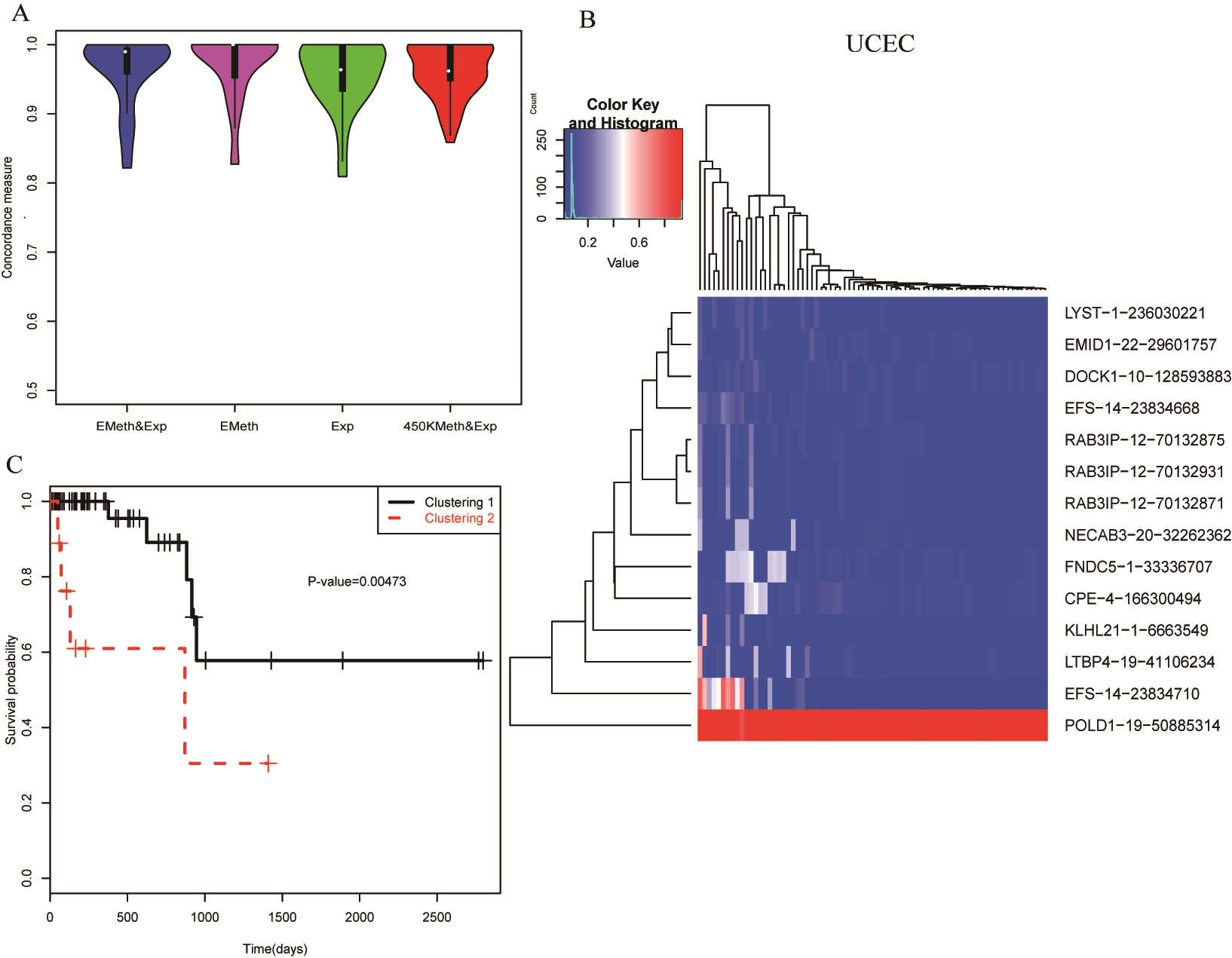

Figure S19. Survival analysis of UCEC using the features selected in more than 20% of the cross validations. Each row is a selected feature (gene expression or methylation locus) and each column is a tumor sample.

Table S1. The tissue/cell line list used to train the EAGELING model

| WGBS ID | 450K ID | Tissue/cell line name |
| --- | --- | --- |
| GSM675542 | GSM1139417 | H1 cell line |
| GSM432687 | GSM1139405 | IMR90 cell line |
| GSM916049 | GSM1270017 | liver cell |
| GSM983647 | GSM1281096 | Lung Cell |
| GSM1172596 | GSM1270015 | fetal leg muscle tissue |
| GSM1120326 | GSM1129708 | Aorta tissue |
| GSM983651 | GSM1281098 | Pancreas tissue |
| GSM916050 | GSM1546443 | Hippocampus Middle |
| GSM706059 | GSM1139404 | H9 cell line |
| GSM1010983 | GSM1500871 | Fat Cell |
| GSM1010981 | GSM868080 | Adrenal Cells |
| GSM983649 | GSM1276756 | Esophagus cell |
| GSM983646 | GSM1219869 | Small Bowel Cell |
| GSM983652 | GSM868069 | Spleen cell |
| GSM1172595 | GSM868064 | Fetal Thymus cell |
| NIC1254A95 | BLCA-BT-A20V-11A-11D-A14Z-05 | Normal bladder sample |
| A7-A0CE-11A-21D-A148-05 | BRCA-A7-A0CE-11A-21D-A10Q-05 | Normal breast sample |
| NIC1254A70 | LUAD-44-6148-11A-01D-1756-05 | Normal lung sample |
| NIC1254A75 | STAD-BR-6452-11A-01D-1801-05 | Normal stomach sample |
| NIC1254A89 | BLCA-BL-A13J-01A-11D-A276-05 | Bladder urothelial carcinoma sample |
| NIC1254A93 | BLCA-BT-A2LA-01A-11D-A18G-05 | Bladder urothelial carcinoma sample |
| NIC1254A91 | BLCA-BT-A20V-01A-11D-A14Z-05 | Bladder urothelial carcinoma sample |
| NIC1254A96 | BLCA-DK-A1AA-01A-11D-A13Z-05 | Bladder urothelial carcinoma sample |
| NIC1254A94 | BLCA-DK-A1AG-01A-11D-A13Z-05 | Bladder urothelial carcinoma sample |
| NIC1254A92 | BLCA-H4-A2HQ-01A-11D-A17Y-05 | Bladder urothelial carcinoma sample |
| NIC1254A17 | BRCA-A2-A0YG-01A-21D-A10A-05 | Breast invasive carcinoma sample |
| NIC1254A69 | LUAD-44-6148-01A-11D-1756-05 | Lung adenocarcinoma sample |
| NIC1254A71 | LUAD-67-6215-01A-11D-1756-05 | Lung adenocarcinoma sample |
| NIC1254A110 | STAD-BR-6452-01A-12D-1801-05 | Stomach adenocarcinoma sample |
| NIC1254A76 | STAD-CG-5730-01A-11D-1601-05 | Stomach adenocarcinoma sample |
| NIC1254A77 | STAD-D7-6519-01A-11D-1801-05 | Stomach adenocarcinoma sample |
| NIC1254A78 | STAD-F1-6177-01A-11D-1801-05 | Stomach adenocarcinoma sample |
| NIC1254A111 | UCEC-AX-A1CK-01A-11D-A138-05 | Uterine corpus endometrial carcinoma sample |

Table S2. The 5 triple-evidenced genes most commonly shared among the 13 cancers

| Gene name | #cancers | Cancer type |
| --- | --- | --- |
| CELSR3 | 11 | BLCA,BRCA,COAD,ESCA,KIRC,KIRP,LUAD,LUSC,PRAD,HNSC,UCEC |
| TNXB | 11 | BRCA,COAD,ESCA,KIRC,KIRP,LIHC,LUAD,LUSC,THCA,HNSC,UCEC |
| TRPM2 | 11 | BLCA,BRCA,COAD,KIRC,KIRP,LUAD,LUSC,PRAD,THCA,HNSC,UCEC |
| KCNAB1 | 10 | BLCA,BRCA,COAD,ESCA,KIRC,LUAD,LUSC,PRAD,THCA,UCEC |
| TRIP13 | 10 | BRCA,COAD,KIRC,KIRP,LIHC,LUAD,LUSC,PRAD,HNSC,UCEC |

Table S3. The top 5 triple-evidenced genes that could not be identified using the original 450K methylation array

| Triple-evidenced gene name | #Type of cancers | Cancer type |
| --- | --- | --- |
| FANCI | 8 | BLCA,BRCA,KIRC,KIRP,LIHC,LUAD,THCA,UCEC |
| RECQL4 | 8 | BLCA,BRCA,COAD,KIRP,LIHC,LUSC,HNSC,UCEC |
| TACC3 | 8 | BLCA,COAD,ESCA,KIRC,LIHC,LUSC,HNSC,UCEC |
| CLU | 7 | BRCA,COAD,KIRP,LUAD,LUSC,HNSC,UCEC |
| SIK1 | 6 | BRCA,KIRP,LIHC,LUAD,LUSC,UCEC |

Table S4. Predictive features selected in more than half of the cross validations in the pan-cancer diagnosis. For the methylation features, they are notated as “Gene name-chromosome number-genomic coordinate”.

A2M, MMP9, TOP2A, LOXL2, CTHRC1, MYL9, CX3CL1, FRZB, LAMA4, MMRN2, PCDH12, PDE2A, RCSD1, TNXB-6-32015635, TNXB-6-32015841, RRM2-2-10261055, CELSR3-3-48701964, CELSR3-3-48701999, SLC16A3-17-80187971, SLC16A3-17-80186222, FANCI-15-89787018, MMP11-22-24115386, MMP11-22-24115407, SIK1-21-44848478, SIK1-21-44848775, TRIM59-3-160167977, PRX-19-40919245, LOXL2-8-23262159, MSH5-6-31706315, CDCA5-11-64853916, PAQR4-16-3018155, HOXD8-2-176992659, LILRB1-19-55141424, CBX7-22-39548103, DNMT3B-20-31366149, NYNRIN-14-24867491, ACVRL1-12-52301580, CDH5-16-66400535, COL5A2-2-190044983, FLNC-7-128469922, GAS7-17-9930882, KRT80-12-52587137, LOXL2-8-23262023, PAMR1-11-35547925, PAQR4-16-3018752, PRX-19-40918880, RCAN1-21-35897562

Table S5. The top 20 features selected in the cross validations of discriminating individual cancers (multi-class classification). For the methylation features, they are notated as “Gene name-chromosome number-genomic coordinate”.

NFIX, SUSD2, PC, CLU-8-27472395, MITF-3-69812973, CYGB, BOC, LAMA4-6-112575929, MRVI1, CRYAB, ABLIM3, CDCA5, LAMB3, ERG-21-40033590, PER3-1-7843618, CD248-11-66084197, AHNAK, SLC7A8, C3, HMGA1-6-34204440

Table S6. Features important in the prognosis analysis. For the methylation features, they are notated as “Gene name-chromosome number-genomic coordinate”.

CELSR3, C2, SLC12A8, TNXB-6-3268893, RRM2-2-10262434, ALS2CL-3-46734833, C2-6-3162871, CELSR3-3-48700375, DBNDD1-16-90077956, LTBP4-19-41107077, SLC12A8-3-124931590, SLC15A3-11-60720297, SLC16A3-17-80190195, ADORA2A-22-24819797, AHNAK-11-62314602, AMOTL2-3-134093532, BMP1-8-22021601, CYR61-1-86045945, FANCI-15-89787087, HLA-A-6-1419731, HLA-DRA-6-32406271, ITGB4-17-73720591, KCTD12-13-77461185, KIF22-16-29801788, LAMA3-18-21269410, LEPR-1-65886336, LINGO1-15-78114220, MECOM-3-169382163, MICAL2-11-12159354, MICB-6-2750441, MMP11-22-24115466, MYH11-16-15952247, MYLK-3-123602919, PFKFB3-10-6186637, RGMA-15-93632389, RORA-15-61521887, SIK1-21-44846688, SLC17A9-20-61582071, SMARCD3-7-150974361, THRB-3-24535866, TMC4-19-54677337, TMEM132A-11-60690872, TRIM2-4-154072288, TRIM59-3-160167317, UHRF1-19-4909443, WDR90-16-699218

Table S7. Sample size of the 13 cancers in the TCGA database

| Cancer type | DNA methylation | | Gene expression | |
| --- | --- | --- | --- | --- |
|  | Normal | Tumor | Normal | Tumor |
| BLCA | 21 | 419 | 19 | 408 |
| BRCA | 97 | 797 | 113 | 1102 |
| COAD | 38 | 314 | 41 | 287 |
| ESCA | 16 | 186 | 11 | 185 |
| HNSC | 50 | 530 | 44 | 522 |
| KIRC | 160 | 325 | 72 | 534 |
| KIRP | 45 | 276 | 32 | 291 |
| LIHC | 50 | 380 | 50 | 373 |
| LUAD | 32 | 475 | 59 | 517 |
| LUSC | 42 | 370 | 51 | 501 |
| PRAD | 50 | 503 | 52 | 498 |
| THCA | 56 | 515 | 59 | 513 |
| UCEC | 46 | 439 | 24 | 177 |

Table S8. The sample sizes of different cancers in the survival analysis

| Cancer | Total | Living | deceased |
| --- | --- | --- | --- |
| BLCA | 111 | 81 | 30 |
| BRCA | 807 | 710 | 97 |
| COAD | 384 | 329 | 55 |
| ESCA | 162 | 105 | 57 |
| HNSC | 314 | 190 | 124 |
| KIRC | 458 | 312 | 146 |
| KIRP | 100 | 86 | 14 |
| LIHC | 59 | 33 | 26 |
| LUAD | 310 | 227 | 83 |
| LUSC | 319 | 198 | 121 |
| UCEC | 450 | 409 | 41 |
